# Heterogeneity in collective endothelial cell behavior is a driver of arterio-venous remodeling

**DOI:** 10.1101/2020.09.15.297093

**Authors:** Keyi Jiang, Cathy Pichol-Thievend, Zoltan Neufeld, Mathias Francois

## Abstract

During vascular development, arteries and veins form in a stepwise process that combines vasculogenesis and sprouting angiogenesis. Despite extensive data on the mechanisms governing blood vessel assembly at the single cell level, little is known about how cell populations migrate in a finely tuned and coordinated manner, and distribute precisely between arteries and veins. Here, we use an endothelial-specific zebrafish reporter, *arteriobow*, to label small cohorts of arterial cells and trace their progeny from the initial events of vasculogenesis through the process of arterio-venous remodeling. We reveal that the genesis of both arteries and veins relies on the coordination of ten types of collective cell behaviors originating from discrete endothelial cell clusters. Within these behavioral categories, we identify a heterogeneity of collective cell dynamics specific to either arterial or venous remodeling. Using pharmacological blockade, we further show that factors known to control vascular patterning such as cell-intrinsic Notch signaling and cell-extrinsic blood flow, potentially act as regulators by coordinating endothelial cohorts behavior, which in turn instructs the future territory of arterio-venous remodeling.

## Introduction

A functional blood vasculature is essential for physiological homeostasis, through efficient transport and supply of oxygen, nutrients, metabolites and cells to almost all tissue types. The vertebrate vascular system initially develops as one primitive network, which then undergoes remodeling to give rise to distinct vessel subtypes including arteries and veins (Hogan and Schulte-Merker 2017, Potente and Mäkinen 2017). The molecular basis of blood vessel assembly has been studied for decades, primarily by forward and reverse genetic approaches, leading to the identification of key signaling pathways that are essential for endothelial cell specification and vascular patterning. Endothelial cells (ECs) form the inner lining of all blood vessels. While sharing a mesodermal origin, EC populations of various vascular beds have the potential to give rise to distinct specialized subtypes (Aird 2012, Potente and Mäkinen 2017, Pichol-Thievend, Betterman et al. 2018). These distinct sub-populations display a wide range of individual and collective behaviors that contribute to the establishment of organ-specific networks. Characterization of EC assembly demonstrates that blood vessel morphogenesis is a complex process involving the spatiotemporal coordination of a collection of cell behaviors (Wacker and Gerhardt 2011, Jianxin, Castranova et al. 2015). This includes change in cell polarity, migration, proliferation, apoptosis, differentiation and variability in cell-to-cell contacts (Jakobsson, Franco et al. 2010, Geudens and Gerhardt 2011). Further, ECs have a remarkable plasticity and exhibit heterogeneous morphological and molecular signatures to meet specific needs of the vessels and organs in which they reside (Torres-Vázquez, Kamei et al. 2003, Aird 2012, Potente and Mäkinen 2017).

The zebrafish is the *in vivo* model of choice to study vascular morphogenesis in real time at cellular resolution, due to superior live imaging technology, combined with rapid genome editing and transgenesis capabilities (Hogan and Schulte-Merker 2017). In the trunk of the zebrafish embryo, vasculogenesis originates from mesodermal progenitor cells in the lateral plate mesoderm that specialize into angioblasts at around 12 hours post fertilization (hpf) (Zhong, Rosenberg et al. 2000, Sumanas and Lin 2005). At 14-20 hpf, these angioblasts further differentiate into arterial and venous endothelial cells (ECs) to form two major axial vessels, the dorsal aorta (DA) and posterior cardinal vein (PCV) (Kohli, Schumacher et al. 2013). Following assembly of these main axial vessels, sprouting angiogenesis commences at around 22 hpf, with ECs sprouting from the DA in between the somites to form primary ISVs (intersegmental vessels). Leading ECs in adjacent sprouts subsequently join one another on the dorsal side of the trunk to further form the dorsal longitudinal anastomotic vessel (DLAV) (Isogai, Lawson et al. 2003). This process whereby ECs assemble to form the ISV and DLAV implies that the primitive vasculature is exclusively arterial and that, through a series of vascular remodeling events, a fraction of the primary vasculature acquires a venous identity (Bussmann, Bos et al. 2010). The morphological process that ultimately organizes the trunk blood vascular system into an exquisitely balanced ratio of 1:1 arteries (aISVs) and veins (vISVs), begins at around 30-32 hpf when a second wave of sprouting vessels emerges from the PCV (Isogai, Lawson et al. 2003, Küchler, Gjini et al. 2006, Yaniv, Isogai et al. 2006). These venous sprouts either anastomose with adjacent primary ISVs to form a transient three-way connection between the DA, an ISV and the PCV, or migrate dorsally to the horizontal myoseptum (HM) to give rise to paralymphangioblasts (PLCs) which will contribute to the lymphatic vasculature (Isogai, Lawson et al. 2003, Yaniv, Isogai et al. 2006, Geudens, Herpers et al. 2010). In the three-way connected vessels, ISVs either remain stably connected to the DA, hence maintaining an arterial identity, or regress, thus turning the primary ISVs into a vein. These newly formed venous vessels maintain a connection with the arterial network to establish a fully functional blood vascular circuitry.

A number of cellular mechanisms have been proposed to control the precise patterning and distribution of arteries and veins. A recent study suggested that venous identity is established by displacing arterial ECs, with venous ECs migrating from the PCV into primary ISVs without any requirement for endothelial trans-differentiation (Weijts, Gutierrez et al. 2018). Another study showed that ECs are pre-specified early (16-18 hpf) in the lateral plate mesoderm and that ISVs are pre-destined to become either arterial or venous. In this model, EC fate is specified by the regulated activity of local Notch signaling in response to blood flow (Geudens, Coxam et al. 2019). In future aISVs, EC ventral polarity, ventral migration, as well as a multicellular lining at their base are mediated by local constitutively activated Notch signaling. It is thought that this high Notch signaling activity prevents these ISVs from transforming into vISVs in the presence of a transient three-way connection. By contrast, in response to low shear-stress blood flow, low Notch activity in future vISVs promotes EC dorsal polarization and dorsal migration. This results in a change of endothelial cell identity, which will support the formation of future veins (Geudens, Coxam et al. 2019). Despite our understanding of these early specification events, their relationship to the collective cell dynamics that governs the distribution of vessel sub-types remains elusive.

Here, taking advantage of high-resolution confocal live imaging, cell fate mapping with multi-color labeling, and unbiased clustering analysis based on color-distance measurement, we investigate the heterogeneity and cell fate trajectory of the primitive arterial vasculature. Long term tracing (10days) of cell cohorts distribution reveals that unlike the developing muscle tissue whereby an overly dominant clone will establish most of the cell population (Nguyen, Gurevich et al. 2017), the blood vasculature remains extremely heterogenous in nature with a balanced distribution of a variety of cohesive EC clusters. Tracking of distinct groups of physically related ECs over the first 3 days of development demonstrates that arterio-venous remodeling involves an interplay between highly directed mobility, dynamic rearrangement, and apoptosis of arterial endothelial clones. Quantification of collective EC dynamics, at timepoints where the primitive vasculature gives rise to arteries and veins, reveals that arterial ECs display up to ten discrete types of collective behaviors. In particular, our data suggest that heterogeneity in directional migration and spatial re-arrangement of EC clusters predicts future territories of both venous and arterial vascular beds. Further, through manipulating cell-intrinsic Notch activity and cell-extrinsic blood flow, our results provide a new framework for understanding of how discrete clusters of ECs coordinate one another to instruct the proper remodeling of arteries and veins in primary ISVs.

## Results

### Generation of arteriobow reporter transgenic lines

To gain insights into how EC populations organize themselves during blood vessel formation, we monitored how single EC behavior and collective EC dynamics are orchestrated in the early events of vascular patterning. This was achieved through the establishment of an endothelial-specific Brainbow reporter zebrafish line (*fli1:Gal4;10×UAS:Brainbow;fli1:Cre*, Figure 1A). The Brainbow cassette (Livet, Weissman et al. 2007) contains a promoter followed by three spectrally distinct fluorescent proteins: dTomato, mCerulean, and eYFP. For each copy of the construct, only one of these three fluorescent proteins can be expressed, due to the use of two pairs of mutually incompatible *loxp* sites that ensure that Cre-mediated recombination occurs once and is irreversible. The default spectral expression is red, but after recombination, expression is switched to either cyan or yellow (Livet, Weissman et al. 2007, Pan, Livet et al. 2011). In addition, increase in color diversity is directly proportional to the number of copies of the Brainbow transgenes inserted into the fish genome, owing to the stochastic recombination and combinatorial expression of fluorescent proteins. In this study, we kept the number of transgenes to three copies across the different breeding pairs in order to generate a consistent range of color diversity (supp. Figure 1 to Figure 1).

**Figure 1.**
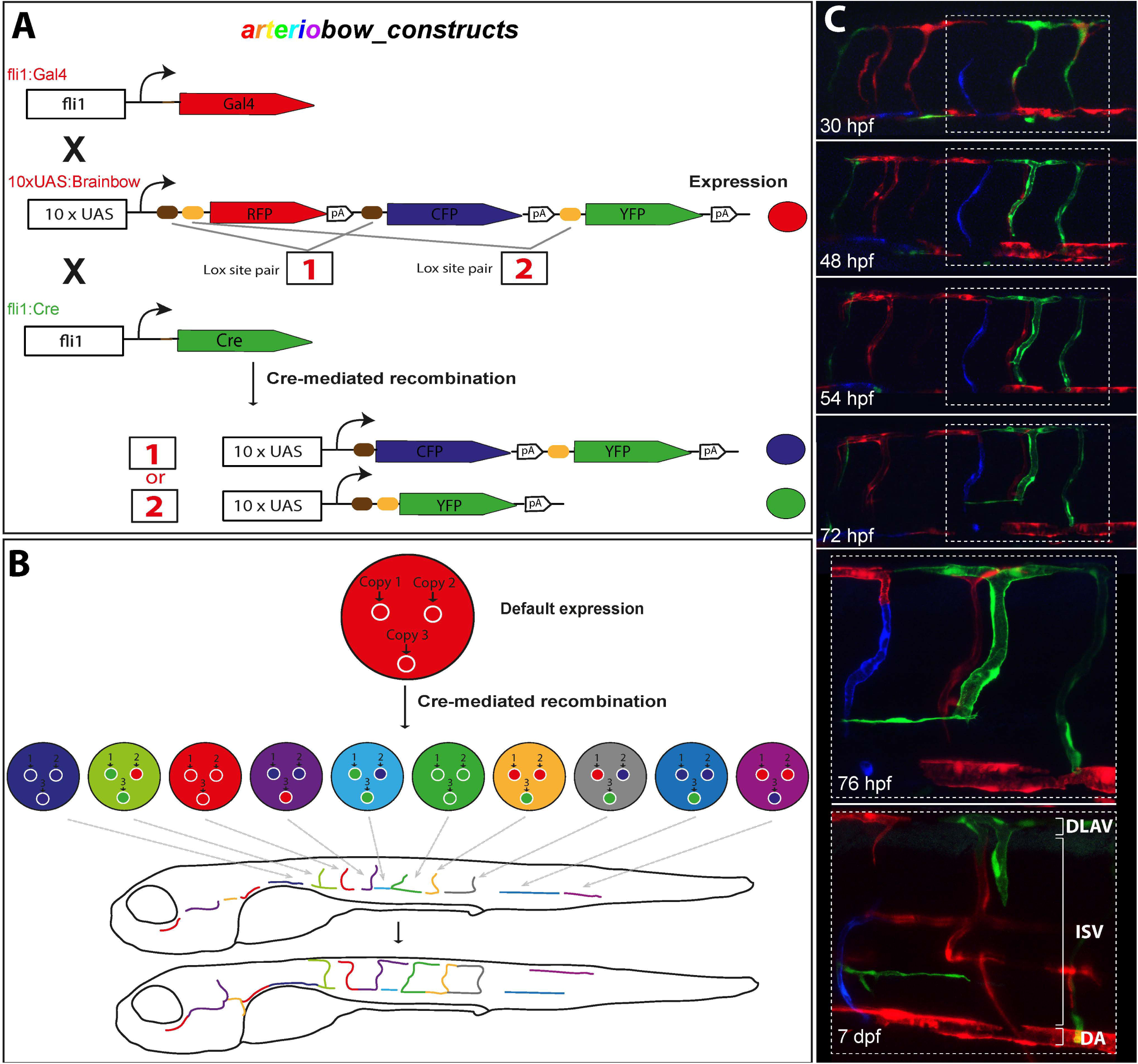
Labelling individual EC cohorts of the blood vasculature using random fluorophore combination. (A) Diagram of the *fli1:Cre;fli1:Gal4*;*10×UAS;Brainbow* (*arteriobow*) reporter system. Transgenic reporter lines were generated using the *fli1:Gal4, 10XUAS:Brainbow*, and *fli1:Cre* constructs. Each copy of the Brainbow cassette comprises three spectrally different fluorescent proteins and two pairs of mutually exclusive Loxp sites. The Cre recombinase excises only once either pair 1 (brown box) or 2 (yellow box). (B) Spectral colour diversity derived from *arteriobow* illustrated for single cells and trunk vasculature with three copies of the Brainbow construct. Each insert is recombined randomly with three possible outcomes (red, blue or green), resulting in the production of ten possible combinations. (C) Different representative time points of *arteriobow*, which has two copies of the reporter cassettes. Insets at 76 hpf and 7 dpf are close ups of the framed areas at 30 hpf, 48 hpf, 54 hpf and 72 hpf. hpf, hours post fertilization; aISV, arterial intersegmental vessel; DA, dorsal aorta; DLAV, dorsal longitudinal anastomosing vessel.

Since a daughter cell will inherit a unique color from its mother cell, this multi-color labeling technology enables sparse labeling of distinct endothelial cell clusters and the tracing of their progeny as vascular development proceeds (Figure 1B). The Brainbow expression is under the control of a Gal4-inducible system, with the presence of 10× non-repetitive UAS sequences (10× UAS:Brainbow, Figure 1A). To achieve endothelial-specific multicolor labeling, we generated *fli1:Gal4* driver line and *fli1:Cre* line (Figure 1A). The *fli1* gene belongs to the Ets transcription factor family and is one of the earliest transcription factors driving both blood and endothelial development (Liu, Walmsley et al. 2008, Park, Kim et al. 2013). The use of the *fli1* promoter in *fli1:Gal4;UAS:Brainbow*;*fli1:Cre* enabled us to mark early endothelial progenitors with a unique combination of fluorophores. It is important to note that the transgenic reporter lines *fli1:Gal4;UAS:Brainbow* were generated as mosaics and that those selected for inclusion in this study display a bias towards labeling almost exclusively arterial EC populations (Figure 1C). The reason *fli1* transgene activity was restricted primarily to certain arterial sub-populations remains elusive, but it could be due to mosaic expression. This is a commonly seen phenomenon with the use of the Gal4/UAS system, resulting in only a subset of arterial progenitor cells being labeled in some transgenic founders (Scott, Mason et al. 2007, Asakawa and Kawakami 2008). Regardless, this bias was advantageous in the context of this study to enable preferential tracking of arterial cell populations. As a consequence, we named this Brainbow reporter system “*arteriobow*”.

### A dynamic repertoire of collective EC behaviours during vascular remodelling

To study the behavioral dynamics of discrete arterial EC clusters, we performed lineage tracing at the same anatomical location across different time points during embryonic development. Specifically, we imaged the posterior yolk extension area comprising an average of 5 to 7 inter-somitic vessels, using the very posterior end of the yolk extension as an anatomical reference (Figure 2A). In order to assign ECs to a specific cluster and establish their lineage relationship, we developed a method to measure the color distance within and between different groups of cells in an unbiased manner (supp. Figure 1 to Figure 2). We tracked arterial EC clusters from 30 hpf to 72 hpf. We found that ECs form cohesive cohorts, mobilize as a collective, and exhibit a variety of collective behaviors. Of note, in rare instances small grouping of cells become disjointed and move independently from cells with the same lineage. Through examining a large number of individual EC collectives, we identified, classified and quantified ten types of collective cell behaviors (Figure 2B-D). Type I (9%), ECs migrate dorsally from the DA into the ISV, but only over a short distance (∼73 uM on average). Once into the ISV, these cells either remain static or migrate back, slightly ventrally, towards the DA (All described types I-X are displayed on Figure 2C-C’’’). Type II (5%), endothelial clusters migrate dorsally along the ISV and stop their migration upon contact with the DLAV. Type III (11%), groups of cells migrate from the ISV further into the DLAV, forming a “T” shape structure connecting the two vessel subtypes. Type IV (17%), endothelial clusters form a transient “T” shape rapidly changing into an “L” form structure bridging the ISV and DLAV. Type V (12%), EC clusters migrate out from the ISV into the DLAV and assemble as a line segment. Type VI (4%), cell clusters split into two and cells migrate in opposite directions in the DLAV. Type VII (14%), groups of endothelial cells rearrange in the DLAV and migrate ventrally into either an ipsilateral or contralateral adjacent ISV. Type VIII (14%), endothelial clusters revert their migration path after reaching the DLAV, migrating back ventrally into their ISV of origin. The Type VIII still occupies the entire ISV. Type IX (5%), by contrast, only cover half of the ISV while showing similar series of migrating events to the type VIII. Type X (11%), ECs undergo apoptosis during vascular pruning.

**Figure 2.**
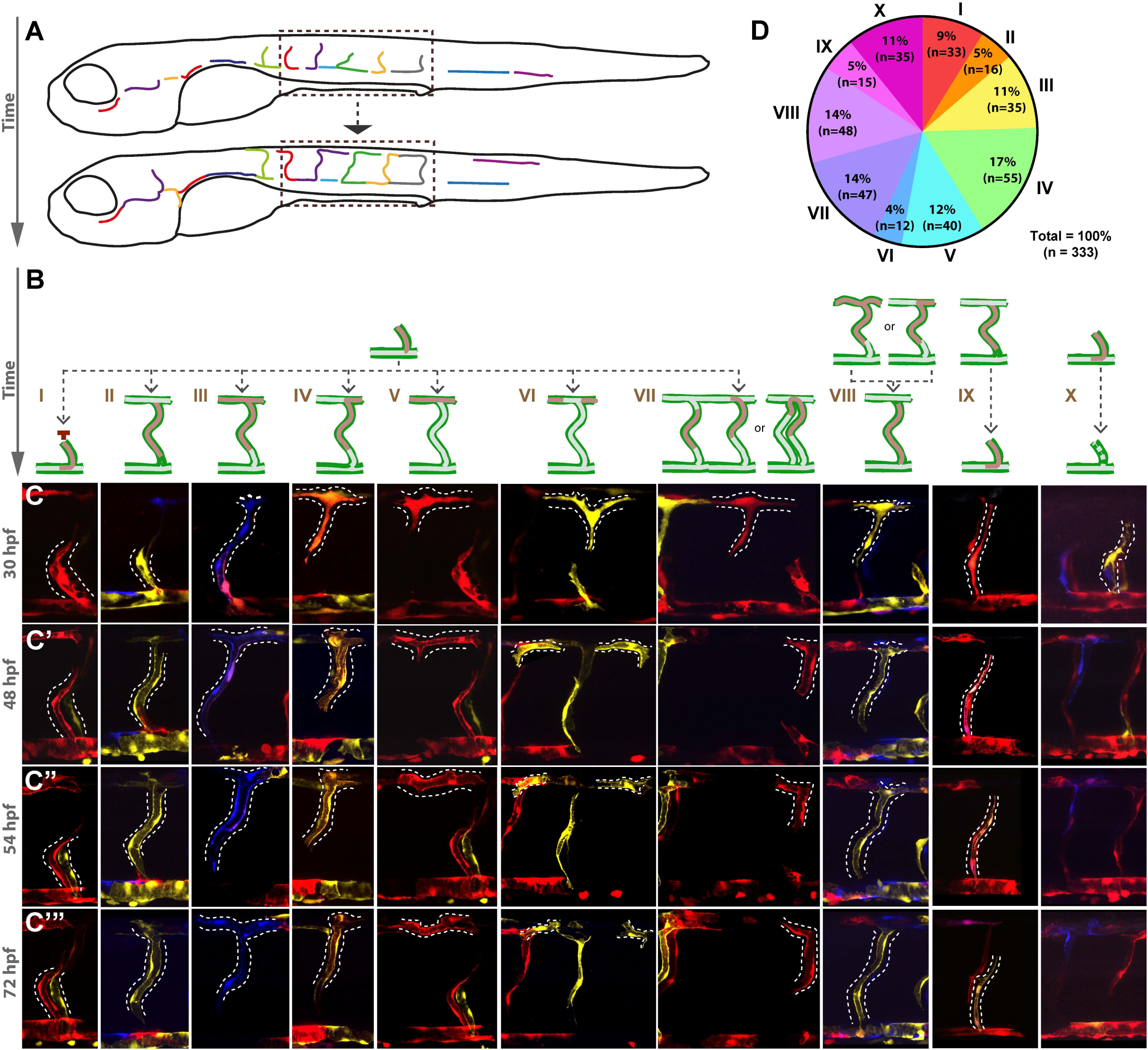
Heterogenous behaviors of endothelial cell cohorts contribute to ISV/DLAV remodeling. (**A**) Schematic of EC cluster tracing during vascular remodeling. The analysis was consistently performed in the yolk sac extension over 8-10 somites. (**B**) Morphological classification of cohesive EC clusters over time reveals ten classes of collective EC behaviors. (C-C’’’) Four representative time points of EC cluster tracing over vascular remodeling, 30 hpf (C), 48 hpf (C’), 54 hpf (C”), and 72 hpf (C’”), respectively. (D) Quantification of the proportion of EC clusters found in each of the 10 behavior classes. n=6 experiments, 42 embryos, 333 clusters. hpf, hours post fertilization.

Further, we performed a longer-term lineage tracing analysis from 24 hpf to 10 dpf to assess whether the initial heterogeneity is preserved as the vascular network matures. The heterogeneity across the whole trunk was analyzed in an unbiased and high throughput manner (number of fish = 6, number of cells per fish ∼ 200, see method principle in supp. Figure 2 to Figure 2). A striking feature of the 10-dpf vasculature is that the pattern of color distribution closely resembles that observed at 24 hpf. This suggests that the original heterogeneity acquired in the initial events of vessel morphogenesis is not lost to a subset of super-dominant endothelial cohorts, as has been observed during muscle development (Nguyen, Gurevich et al. 2017).

### Collective behaviour of EC cohort indicates future territories of arterial or venous remodelling

There is compelling evidence that vascular remodeling involves various EC events, including directed EC movement (Chen, Jiang et al. 2012, Udan, Vadakkan et al. 2013, Franco, Jones et al. 2015, Rochon, Menon et al. 2016, Geudens, Coxam et al. 2019), rearrangement (Xu, Hasan et al. 2014, Franco, Jones et al. 2015), proliferation (Xu, Hasan et al. 2014, Fang, Coon et al. 2017) and death (Hughes and CHAN - LING 2000, Lobov, Rao et al. 2005). Recently, directed migration of ECs has been shown to be correlated to arterio-venous remodeling (Weijts, Gutierrez et al. 2018, Geudens, Coxam et al. 2019). Here we show that EC cohorts labelled with the same color and therefore derived from the same lineage, migrate directionally (either dorsally or ventrally along ISVs), rearrange (type VII migrating from one ISV into an adjacent ISV) and undergo cell apoptosis (type X). This prompted us to hypothesize that differential behavior of EC collectives could be used as a readout to inform future locations of arterio-venous remodeling. To identify arterial ISV (aISVs) and venous ISVs (vISVs) as well as their connection, respectively, to the DA or the PCV, the *arteriobow* line was crossed into a *kdrl:mCherry* background, a blood vessel reporter system. The behavioral types of EC collectives were characterized and quantified by conducting a long-term lineage tracing analysis from 30hpf to 5dpf. At the end point of the lineage tracing, we further assessed the connection of ISVs to the DA or the PCV and the direction of blood flow in order to assign arterial or venous identity to ISVs (Figure 3).

**Figure 3.**
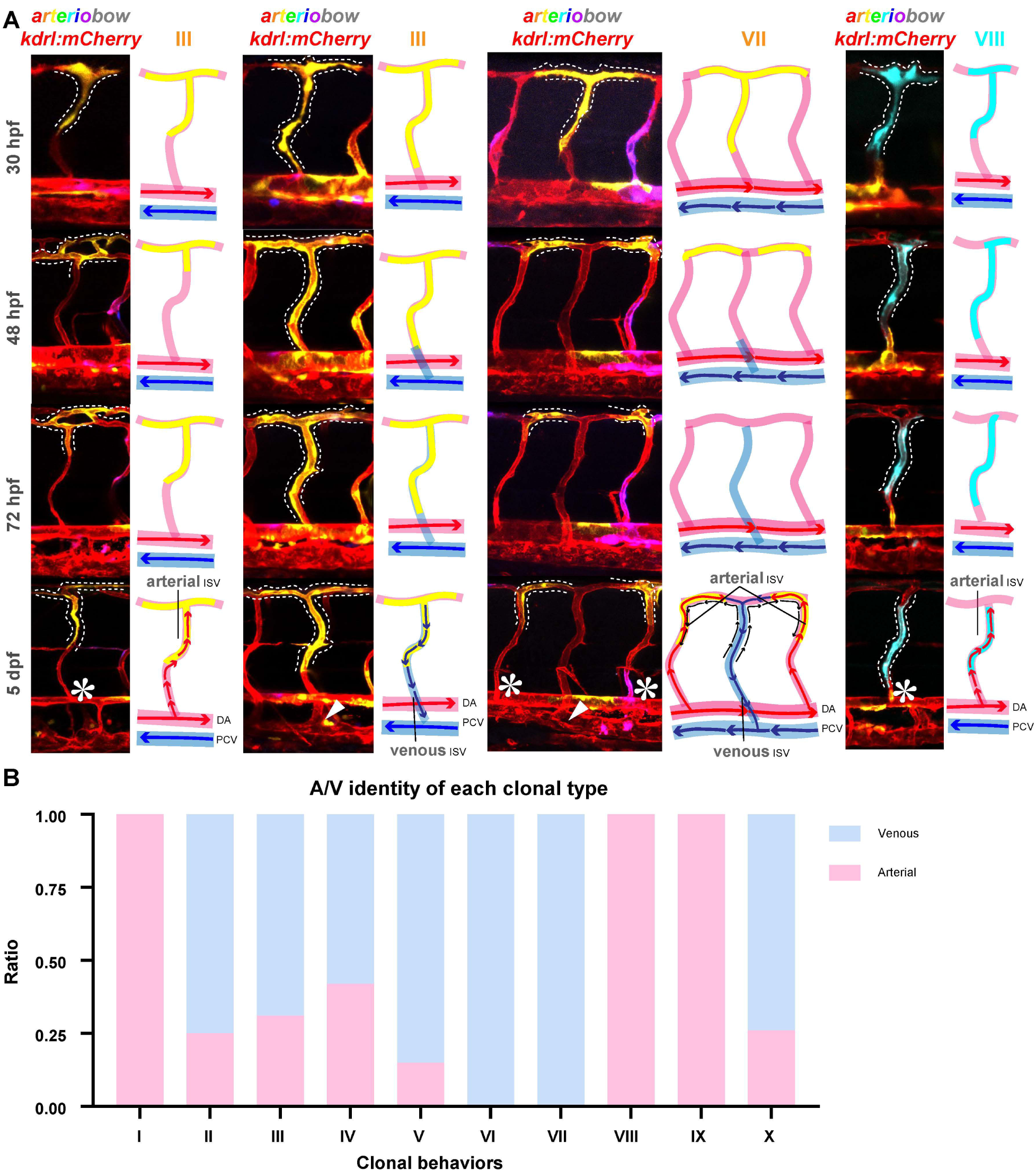
Relationship between collective behaviors of arterial endothelial cells and arterio-venous remodeling.. (**A**)The *arteriobow* line was crossed to the *kdrl:mCherry* line and EC cohorts were analyzed and mapped in relation to arterio-venous specification. A subset of representative collective EC behaviors (III, VII and VIII) are outlined by a dashed line. Asterisks denote the connection between ISVs and the DA, while arrowheads show the connection between ISVs and the PCV. (**B**) Quantification of the ratio of arterial and venous ISVs assigned to a specific type of collective EC behavior. Analysis performed in the yolk extension region over 10 somites from 30 hpf to 5 dpf *arteriobow;kdrl:mCherry* embryos (n=6 experiments, 42 embryos, 333 clusters). hpf, hours post fertilization; dpf, days post fertilization; aISV, arterial intersegmental vessel; DA, dorsal aorta; DLAV, dorsal longitudinal anastomosing vessel.

During primary angiogenesis, all types of endothelial clusters migrate dorsally in a collective manner (Supplementary Video). During vascular remodeling, cell cohorts with type I behavior, if motile, were found to migrate slightly ventrally (supp. Figure 1-2 to Figure 3); types VIII and IX were also found to migrate back along their original migration path in a ventral direction (Figure 3A & supp. Figure 1-2 to Figure 3). When defining ISV identity and correlating it with these three types of behavior (I, VIII, IX), we found that these collective EC activities are exclusively restricted to the arterial compartment (DA/aISV segments, Figure 3B). This observation suggests a correlation between ventral migration of EC collectives and maintenance of their arterial cell identity. The type II, III, IV and V were found to migrate dorsally but be spatially restricted in ISV/DLAV segments, regardless of their venous or arterial identity (Figure 3 & supp. Figure 1-2 to Figure 3). Intriguingly, there was a preferential outcome (60-80%) towards venous formation in ISVs which (originally) contained these endothelial clusters (Figure 3B). As opposed to the ventral migration and subsequent arterial identity characteristic of behavioral types I, VIII and IX, the dorsal migration of types II-V appears to be an indicator of presumptive venous territory. The dorsally migrating type VI and VII are found to change their migration direction in the DLAV: cells from the type VI split and migrate in opposite directions in the DLAV while type VII cells relocate into an adjacent ISV. Interestingly, we observed that the ISVs which originally contained these two cell types (VI and VII) always acquire a venous identity, while the ISVs which type VII cells migrated into remain arterial in nature (Figure 3 & supp. Figure 1-2 to Figure 3). Type X cells were found to undergo cell death during vascular pruning, especially during the resolution of three-way connections among the DA, ISV and the PCV into arterial or venous ISV (supp. Figure 1-2 to Figure 3).

To further validate the role of cell dynamics in determining arterial and venous identity, we measured the proliferation rate and migration distance, which revealed that EC cohorts confined to ventral locations in the DA/aISV segments have a lower proliferation and migratory activity; as compared to types that are more dorsally located (supp. Figure 3 to Figure 3). After migrating out from the DA, EC cohorts fated to remain arterial show relatively smaller cluster size (1-3 cells) and low proliferation rates and a migration over short distances (<150 µm, types I VIII and IX). By contrast larger clusters (4-6 cells) destined to venous territories showed longer migration distances (supp. Figure 3 to Figure 3). Overall, these findings show that heterogeneity between EC cohorts is associated with a wide range of collective cell dynamics throughout the process of arterio-venous remodeling. These results suggest that the activity of discrete EC collectives is highly coordinated at specific locations within the initial vascular plexus, and this activity is tightly correlated with the outcome of arterio-venous remodeling.

### Notch-directed collective migration is a modulator of arterio-venous remodeling

The observation of collective cell movement that is spatially restricted, akin to a tip/stalk cell relationship (Isogai, Lawson et al. 2003, Jakobsson, Franco et al. 2010), prompted us to investigate known genetic pathways that regulate cell-to-cell interaction dynamics. Notch signalling plays an important role both in dictating arterio-venous fate and in the establishment of tip/stalk cell identity (Zhong, Rosenberg et al. 2000, Lawson, Scheer et al. 2001, Zhong, Childs et al. 2001, Chiang, Fritzsche et al. 2017). It has been shown that the chemical inhibition of this pathway leads to an increased formation of venous ISVs (Hogan, Herpers et al. 2009, Geudens, Herpers et al. 2010). Therefore, we quantified the effects of pharmacologically manipulating Notch signalling on the different types of collective EC behaviours defined in this study. At 24 hpf, the *arteriobow; kdrl:mCherry* reporter line was treated with the γLsecretase inhibitor DAPT (NL[NL(3,5-difluorophenacetyl)LLLalanyl]LSLphenylglycine tLbutyl ester), until 54 hpf when the determination of ISVs is nearly complete (supp. Figure 1A to Figure 4). As previously reported (Geudens, Herpers et al. 2010, Weijts, Gutierrez et al. 2018), Notch inhibition resulted in a greater proportion of venous ISVs (Figure 4A), further validating the influence of Notch activity in modulating primary ISV specification.

**Figure 4.**
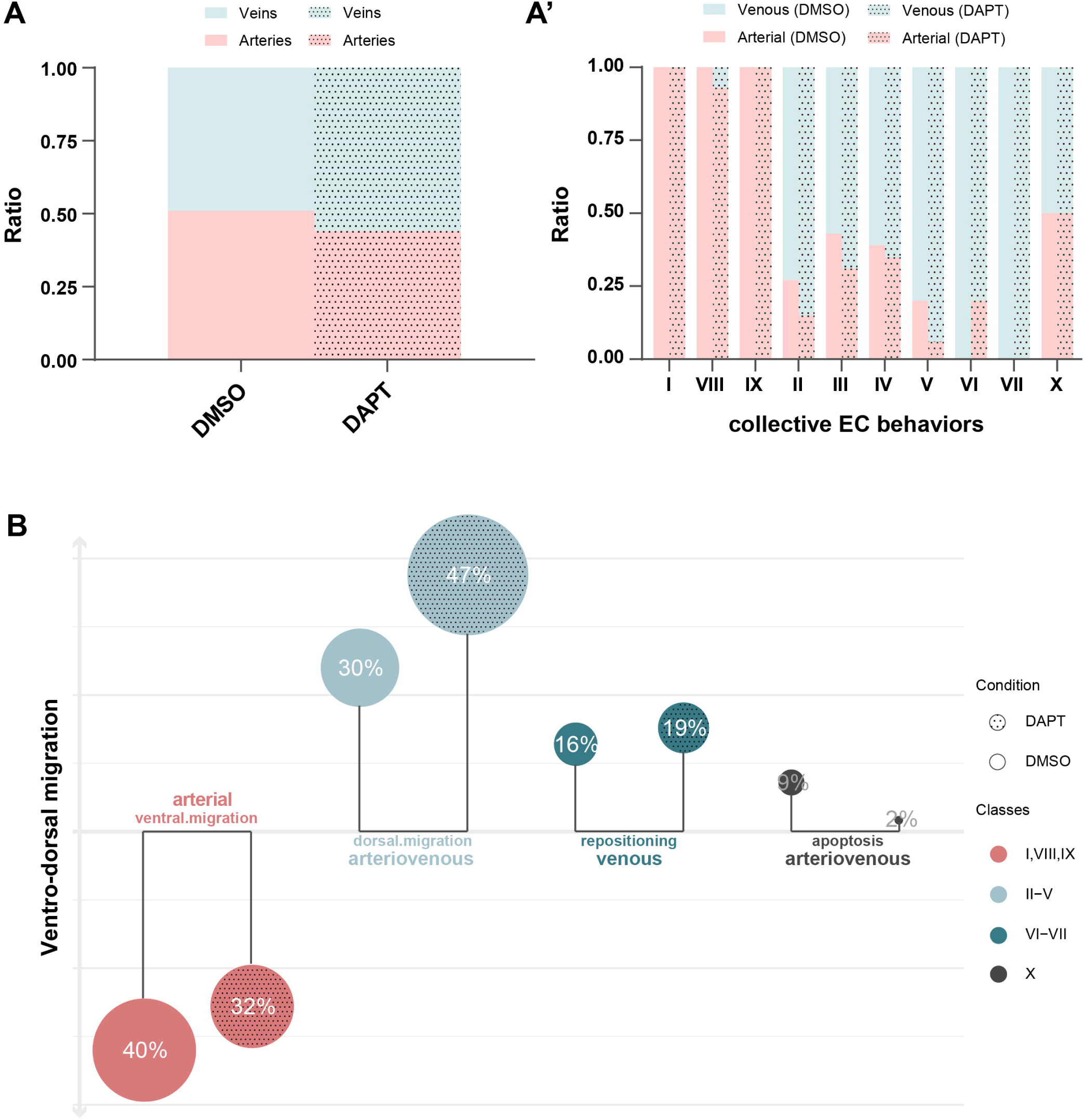
Pharmacological blockade of Notch signaling decreases the proportion of collective behaviors associated with arterial remodeling. **(A)** Quantification of the ratio of arterial and venous ISVs in the yolk extension region comprising ten somites in the trunk of 5 dpf *arteriobow;kdrl:mCherry* embryos either untreated (control; n=3 experiments, 20 embryos) or treated (n=3 experiments, 20 embryos) in presence of DAPT. **(B)** Quantification of the ratio of arterial and venous ISVs assigned to each type of collective EC behavior from the trunk of 5 dpf *arteriobow;kdrl:mCherry* embryos either with or without DAPT treatment. Control (n=3 experiments, 20 embryos, 174 clusters) or DAPT treated (n=3 experiments, 20 embryos, 203 clusters). **(C)** Quantification of the proportion of cohesive EC clusters in each of the four major classes of collective EC behaviors in the *arteriobow;kdrl:mCherry* embryos either treated with or without DAPT. The bottom panel represents the percentage change in ventrally migrating and artery-destined EC collectives (I, VIII, and IX). The top panel shows the percentage change in dorsally migrating and arterio-venous patterning-related EC cohorts (II-V), first dorsally migrating then spatially repositioning and venous remodeling-specific EC clusters (VI-VII), and first dorsally migrating but ultimately dead EC populations (IX).

To monitor the effect of Notch inhibition on the different types of collective EC behaviors, lineage tracing was performed continuously until 5 dpf, when ISV identity is definitive. The frequency of collective behaviors (Figure 1B) was indicative of a venous-dominant outcome (Figure 4A&A’); for instance, dorsally migrating and diverging, and rearranging types showed an increase of 17% (Types II-V) and 3% (Types VI-VII), respectively (Figure 4B). This was paralleled by a 8% reduction in the emergence of arterial-destined behaviors (Figure 4B), including the type I, VIII and IX clusters. The breakdown of decrease or increase by specific behavior is detailed in supp. Figure 1B to figure 4B). These results indicate that a variety of collective EC behaviors, including directional migration, dynamic divergence and rearrangement, are coordinated at a genetic level to shape a global arterial-venous balance. In particular, these results reveal a role for Notch signaling in regulating ISV specification through coordinating distinct behaviors of EC cohorts during arterio-venous remodeling.

### Flow-mediated collective cell motion is essential to arterio-venous remodeling

There is a large body of experimental evidence that vascular remodeling requires blood flow (Le Noble, Moyon et al. 2004, Lucitti, Jones et al. 2007, Chen, Jiang et al. 2012, Udan, Vadakkan et al. 2013). Recent work has uncovered a flow-mediated mechanism that fine-tunes the Notch activity to control EC migratory activity during arterio-venous specification (Weijts, Gutierrez et al. 2018, Geudens, Coxam et al. 2019). Considering the observed effects of Notch inhibition on collective behavior of EC cohorts, we further investigate whether there is a relationship between flow and collective EC behavior. In order to analyze the effects of flow perturbation, the reporter embryos *arteriobow;kdrl:mCherry* were treated with tricaine methanesulfonate (ms-222), which slows down the heart rate and reduces blood flow. The treatment was given from 30 hpf, shortly after the onset of secondary sprouting and prior to anastomosis with the primary (arterial) ISVs, until 54 hpf, when vascular remodeling is near completion (supp. Figure 1A to Figure 5). As previously reported (ref), flow reduction disrupted the overall arterial-venous balance, with a significant increase of 11% in favor of the of the arterial compartment at 5 dpf (Figure 5A).

**Figure 5.**
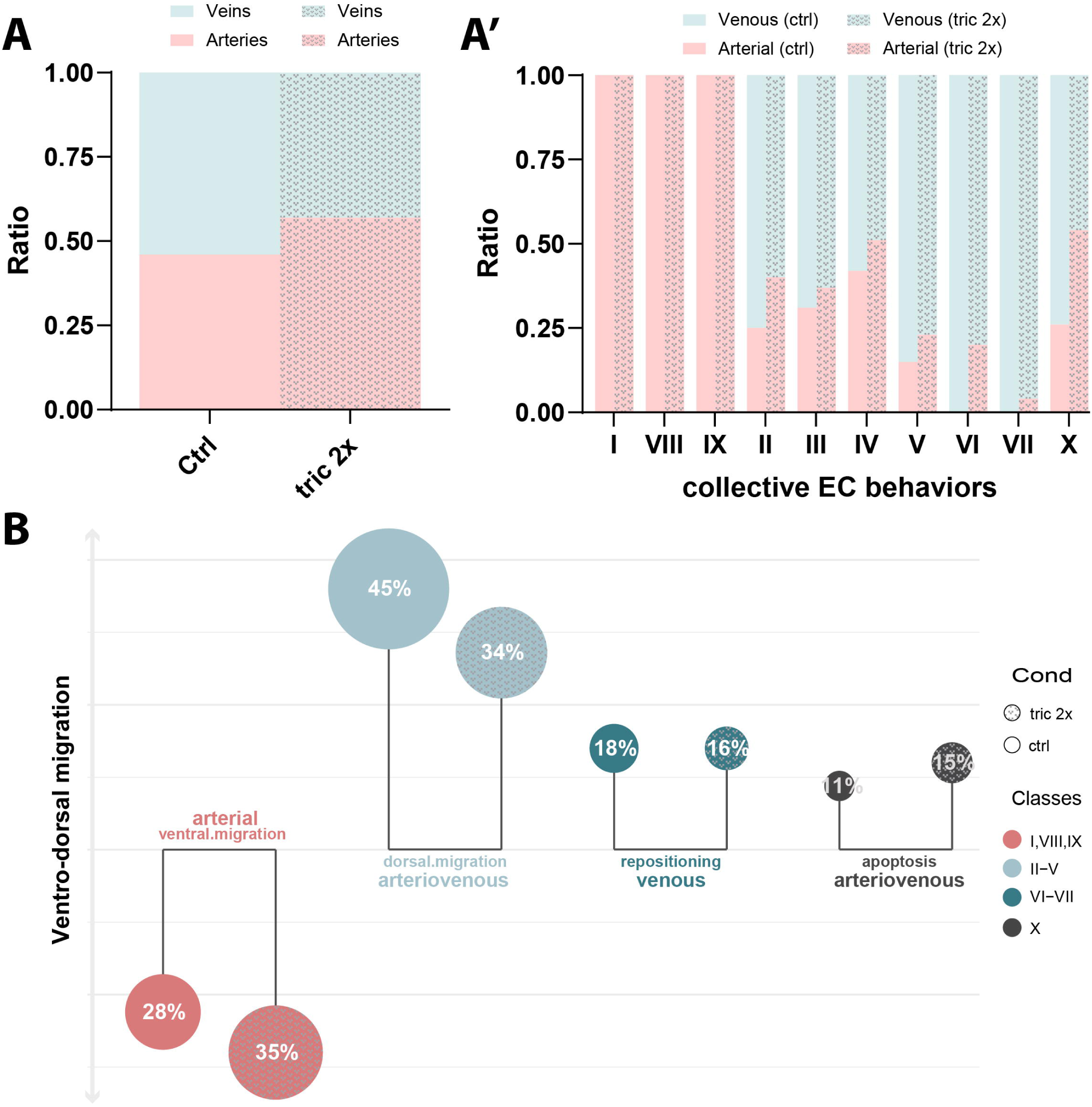
Decrease in blood flow leads to an increase in the proportion of behavioral types associated with arterial remodeling. **(A)** Quantification of the ratio of arterial and venous ISVs in the trunk of 5 dpf *arteriobow;kdrl:mCherry* embryos either untreated (control; n=6 experiments, 42 embryos) or treated (n=6 experiments, 42 embryos) with 2 × tricaine to slow down the blood flow. **(B)** Quantification of the ratio of arterial and venous ISVs assigned to each type of collective EC behavior in the trunk of 6 dpf *arteriobow;kdrl:mCherry* embryos either untreated (control; n=6 experiments, 42 embryos, 333 clusters) or administered with 2 × tricaine (n=6 experiments, 42 embryos, 471 clusters). **(C)** Quantification of the proportion of EC clusters in each of the four major classes of collective behavior**s** was performed in the trunk of 5 dpf *arteriobow;kdrl:mCherry* embryos either in the absence (control; n=6 experiments, 42 embryos, 333 clusters) or presence of tricaine (n=6 experiments, 42 embryos, 471 clusters). The bottom panel represents the percentage change in ventrally migrating and artery-destined EC collectives (I, VIII, and IX). The top panel shows the percentage change in dorsally migrating and arterio-venous patterning-related EC cohorts (II-V), first dorsally migrating then spatially repositioning and venous remodeling-specific EC clusters (VI-VII), and first dorsally migrating but ultimately dead EC populations (IX).

Over the course of 4 days (from 30 hpf to 5 dpf), we investigated collective cell dynamics and correlated collective EC behaviors to the specification of primary ISV arterio-venous identity in both control and tricaine-treated conditions (supp. Figure 1A to Figure 5). In accordance with the overall shift towards more arteries forming upon slowing blood flow (Figure 5A&A’), we found a 7% rise in total number of ventrally migrating EC clusters (Figure 5B). The behavioral types, I, VIII and IX, specific to arteries, increased by 1%, 4% and 2%, respectively (supp. Figure 1B to Figure 5B). Parallel to this expansion of cell behavior correlated to arterial identity, we observed an overall decrease of 11% in the proportion of EC clusters that migrate dorsally and have a bias towards residing in future venous ISVs (Figure 5B). The types III, IV and V, showed a reduction of 3%, 4% and 4%, respectively (supp. Figure 1B to Figure 5B). Further, of the two dorsally migrating types that are specific to venous remodeling, the type VI showed a 2% decrease (supp. Figure 1B to Figure 5B). These results indicate that blood flow acts as an extrinsic cue to coordinate collective EC dynamics, which contributes to a balanced organization of artery and vein formation.

## Discussion

This study shows that heterogeneity in collective EC motion is a key feature of the growing vasculature which enables a proper coordination of arterio-venous remodeling. To gain insights into the complexity of how ECs specialize in a collective fashion, we developed an arterial endothelium-specific reporter tool to observe collective EC behavior and monitor how these dynamics contribute to the process of vascular remodeling (summarized in Figure 6). Through long-term lineage tracing analysis, we observed that discrete EC clusters characterized by collective cell behaviors contribute to the formation of a composite vasculature. These findings are in line with previous observations in the retina and heart vasculature which show that ECs acquire an arterial identity through arresting cell cycle progression (Fang, Coon et al. 2017, Su, Stanley et al. 2018). By contrast, proliferating venous ECs, have been shown to change their migration path over long distances to contribute to the formation of the regenerating fin vasculature (Xu, Hasan et al. 2014).

**Figure 6.**
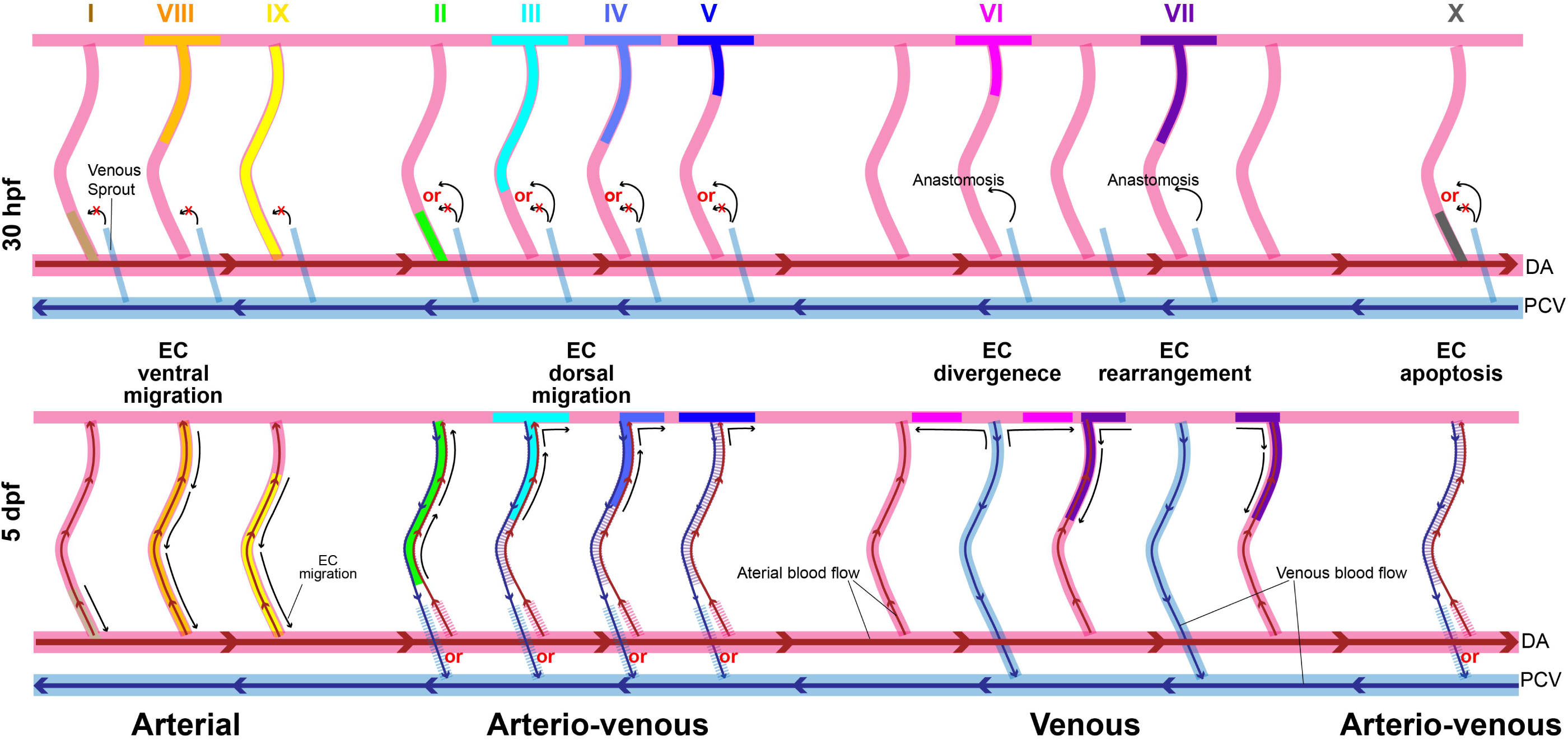
Classification of collective EC behaviors during vascular remodeling. With the use of the Brainbow tool which enables labeling of individual ECs with a unique color and tracking of their progeny, we show that cohesive EC clusters exhibit ten classes of collective behaviors during arterio-venous remodeling, characterized by discrete migratory paths, specific spatial distribution, and characteristic proliferative rate. These behaviors are further categorized into three major collective EC events: (1) directional migration (e.g. ventral and dorsal in the ISVs, and laterally anterior and/or posterior in the DLAV), (2) spatial rearrangement – migrating from the original ISV (future vISV) into adjacent ISV(s) (future aISV(s)), and (3) apoptosis. This substantial heterogeneity in collective EC dynamics is highlighted by behavior variability amongst different EC cohorts which appear to define a signature for future vessel subtype. Specific collective EC behaviors as indicated by numbers are predictive of arterio-venous fate during vascular remodeling. Of importance, behavior VI and VII which predict venous fate are the only two cellular mechanisms whereby EC clusters are disjointed. The balance of profound motility, spatial rearrangement and dynamic apoptosis of endothelial clones is linked to genetic factors and microenvironmental cues. Notch signaling and blood flow synchronize differential collective EC events to drive arterio-venous specification during vascular remodeling. hpf, hours post fertilization; dpf, days post fertilization; aISV, arterial intersegmental vessel; DA, dorsal aorta; DLAV, dorsal longitudinal anastomosing vessel.

Following angiogenesis, vascular remodeling involves vessel differentiation and pruning of an immature vascular network into a perfused and hierarchically branched network. It is increasingly appreciated that collective motion and spatial (re)organization of a physically connected group of cells is one of the fundamental processes dictating tissue patterning during embryonic development (Scarpa and Mayor 2016, Mishra, Campanale et al. 2019, Thüroff, Goychuk et al. 2019). This is supported by our data, demonstrating that collective EC dynamics is tightly related to the arterio-venous remodeling process. Our lineage tracing analysis reveals that vascular remodeling involves a broad diversity of collective EC events: (1) collective directional movement, with EC clusters migrating either in a ventral (type I, VIII and IX) or dorsal (type II-VI) direction, (2) spatial rearrangement, EC collectives (type VII) migrating dorsally into the DLAV from their original ISV, changing migration direction in the DLAV, and migrating ventrally to integrate with the neighboring ISV, and (3) apoptosis, EC cohorts (type X) undergoing cell death.

The collective ventral migration of EC cohorts represented by types I, VIII and IX occurs exclusively in future aISVs, whereas collective dorsal migration of ECs represented by types II-V shows a strong bias towards future vISVs. Migratory behavior of cells in type VI and VII clusters (predictive of venous remodeling) is unique and complex, involving both directional migration and spatial repositioning of ECs. It is possible that directional migration is at play not only during primary angiogenesis (Geudens, Coxam et al. 2019), but also throughout vascular remodeling and may play a major role during the acquisition of arterio-venous identity. Here, we posit that heterogeneity in collective EC movement may drive the acquisition of arterio-venous identity via a co-option of the cell differentiation program by genetic pathways driving EC cohesion and coordinated migration. A possibility is that epigenetic state, transcriptional program and biochemical cues instructed by a collective cell behavior are only permissive to a specific endothelial cell fate. This relationship between collective cell motion and cell fate might be akin to what has been described between cell identity and metabolic state (Tatapudy, Aloisio et al. 2017).

In addition to the role of directional migration and dynamic rearrangement of cohesive EC clusters in arteriovenous differentiation, we note an association of apoptotic events, as indicated by type X clusters, with vessel pruning during vascular remodeling. This largely happens where the intermediate three-way connection between the DA, ISV and the PCV resolves into aISV or vISV. Although it is not yet clear why endothelial cohorts undergo cell death, we hypothesize that this apoptosis might be associated with a failure of ECs to integrate into ISV segments.

This study shows that vessel identity is governed by collective EC dynamics which integrates a coordinated series of events such as velocity, spatial rearrangement, proliferation and apoptosis of endothelial cohorts. During vascular remodeling, the specialization and coordination of ECs is dependent on a wide range of factors, such as cell-intrinsic genetic pathways as well as cell-extrinsic signals from the microenvironment (Herbert and Stainier 2011, Potente and Mäkinen 2017). This work reveals a critical role at play for both the cell-intrinsic Notch signaling pathway and blood flow in coordinating collective EC behaviors during arteriovenous remodeling. Upon Notch perturbation, we show an increase in venous remodeling-associated collective EC behaviors, which includes dorsal migration and spatial (re)organization, paralleled by a decrease in ventral migration events of arterial specific EC cohorts. This change in population dynamics ultimately leads to a patterning of the vasculature biased towards venous remodeling. The acquisition of tip and stalk cell identity during sprouting angiogenesis, relies on a cell at the leading edge (tip cell) which exerts lateral inhibition via the Notch signaling pathway to organize follower cells (stalk cells) (Jakobsson, Franco et al. 2010). Akin to this cell hierarchical assembly, it is possible that the Notch pathway is also required to organize collective EC motion with leaders and followers.

In addition to cell-intrinsic developmental pathways, blood flow is also known to play a critical role in the control of arterio-venous remodeling. Previous reports of directed collective EC migration have shown that vessel pruning is modulated by blood flow (Le Noble, Moyon et al. 2004, Lucitti, Jones et al. 2007, Chen, Jiang et al. 2012, Udan, Vadakkan et al. 2013). Further, venous blood flow-induced dorsal EC migration has been recognized as a mechanism that fine-tunes arterio-venous organization (Weijts, Gutierrez et al. 2018, Geudens, Coxam et al. 2019). In our study, we show that the coordination of directional locomotion of small EC cohorts accounts for a major response to flow in the context of arterio-venous remodeling during vascular development. Decreased blood flow results in increased arterial formation in ISVs, likely due to the diminished effect of flow-mediated compensation on venous remodeling (Geudens, Coxam et al. 2019). In line with this, we show that an increase in ventral migration of arterial-specific EC collectives (I, VIII and IX) is coupled with a decrease in collective behaviors involved in venous remodeling (II-VII), particularly those displaying dorsal migration. We propose that polarized migration of ECs in response to blood flow – dorsal migration against the venous blood flow in future vISV and ventral migration against the arterial blood flow in future aISV – represents an important cellular mechanism to maintain a consistent artery/vein ratio during vascular remodeling. Further, we suggest that mechano-regulation by flow allows the integration of local extrinsic forces to control differential behavior of the EC collectives. For instance, some EC cohorts remain cohesive whereas other EC clusters split and scatter cells around multiple vessel segments. It is therefore likely that blood flow may act as a local organizer of cell to cell coupling which then in turn dictates either parallel (EC collectives maintaining cohesion with unidirectional migration) or swirling motion (EC cohorts splitting and migrating in the opposite direction) (Ladoux and Mège 2017).

In conclusion, our study provides cellular insights into collective EC motion during the assembly of the arterio-venous network. This work uncovers the importance of maintaining heterogeneity in collective EC behavior in order to organize a balanced distribution of arteries and veins. Importantly, of the various factors that are potentially required in collective EC heterogeneity, Notch signaling pathway and blood flow mechanical stimulus appear essential to enable the emergence of and instruct the coordination of this broad diversity of directional migration and spatial repositioning modes of collective EC cohorts to drive arterio-venous remodeling during vascular development. This work opens research directions to further establish a molecular relationship between collective EC behavior and cell fate.

## Methods

### Zebrafish transgenic lines and husbandry

The *Tg(fli1:iCre)* line (referred to as *fli1:Cre*) was generated by a Gateway LR reaction combining p5E-fli1, pME-iCre and p5E-polyA placing the final fli1:iCre sequence into pDestTol2CG2. The *Tg(fli1:Gal4)* line (referred to as *fli1:Gal4*) was kind gift from the Hogan lab (IMB, the University of Queensland). The *Tg(10×UAS:Brainbow)* line (referred to as *UAS:Brainbow*) was generated using the *UAS:Brainbow* construct. *Tg(kdrl:mCherry)* (referred to as *kdrl:mCherry*, labelling blood), *Tg(fli1:nmCherry)* (referred to as *fli1:nmCherry*, labelling all endothelial nuclei) and *Tg(flt1:YFP;lyve1:dsRed)* (referred to as *flt1:YFP;lyve1:dsRed*, flt1 labelling arterial endothelial cells while lyve1 labelling venous endothelial cells) were kindly provided by the Hogan laboratory (IMB, the University of Queensland).

Zebrafish were maintained and bred according to the guidelines of the animal ethics committee at the University of Queensland. Fish embryos were collected from natural crosses and raised at 28.5 0C in E3 embryo medium (5 mM NaCl, 0.17 mM KCl, 0.33 mM MgSO_4_) supplemented with methylene blue (Sigma-Aldrich). At 24 hours post-fertilization (hpf), embryos were transferred to E3 embryo medium with 0.003% 1-phenyl-2-thiourea (PTU; Sigma) to inhibit pigment formation. The embryos were staged as hours or days post fertilization (hpf/dpf) as previously described (Kimmel, Ballard et al. 1995).

### Quantitative PCR for genomic DNA

Genomic DNA was prepared as described by (Dahlem, Hoshijima et al. 2012). Embryos were digested with 50 µl 1M KCl, 0.5M Tris (pH 8), 1M EDTA, 10% IGEPAL, 10% Tween20, MQW, 10 mg/mL proteinase K, incubated at 55°C for 2 hours and then incubated at 99°C for 5 minutes. 1 µl of genomic DNA was mixed with primers and SYBR Green Master Mix (Applied Biosystems) and reactions were run on ViiA7 384 Well Real time PCR with HRM (Bio-Rad). zebrafish genes (*vegfd* and *kdr*) were set as reference, and Brainbow genes (dTomato- and eYFP-coding genes) were used as target genes to represent Brainbow construct. Normalized copy number of Brainbow genes was determined with the comparative Ct method. Copy number of Brainbow construct in the sample = 2 × normalized copy number.

### Live imaging

Embryos were anaesthetized in E3 embryo medium with 0.014% tricaine (MS-222, sigma-Aldrich), mounted in 0.5-1% low melting point agarose with 0.014% Tricaine in 35 mm glass-bottom dishes (20 mm, MatTek), and covered by embryo medium containing 0.004% Tricaine and 0.003% PTU. Confocal images were obtained as z-stacks using Zeiss LSM 710 inverted confocal microscope equipped with temperature (28.5°C) and CO_2_ (5%) controlled chamber. mCerulean, eYFP, and dTomato/mCherry/dsRed were excited by 433 nm, 554 nm, and 574 nm laser lines, respectively.

### Image and color-based clonal analysis

Images were processed with Fiji software (Schindelin, Arganda-Carreras et al. 2012). For clonal analysis, dTomato, eYFP, and mCerulean in figures were rendered in red, green, and blue, respectively. Endothelial cells were identified through identifying their nucleus. Given that the brightest region of an endothelial cell is its nuclear position (supp. Figure 2 to Figure 1), the brightest pixels were unbiasedly selected to represent endothelial nuclei through using the function *find maxima* of Fiji software. Then, the color identity of these selected endothelial nuclei was determined by capitalizing on the basic principle of the Brainbow tool that unique colors were generated by expressing three basic colors, RGB colors, in different ratios in each cell. RGB values, representative of the color identity of the selected endothelial nuclei, were automatically extracted through running a script, “*record maxima colors*”, developed in macro. At last, the clonality of the selected endothelial nuclei was determined based on their color identity. Three methods were developed to assess and visualize their clonality with R software.

*i)* The use of color wheel plots. RGB values of the selected endothelial nuclei were converted to hue and saturation values. On a color wheel, hue represented the color depicted by degrees around the circle, while saturation represented the intensity of the color indicated by the radius. Endothelial cells with similar colors were assumed to have most likely originated from the same parent endothelial cell, and therefore these clonally related endothelial cells were clustered together on color wheels. *ii)* The use of color distance matrices. As R, G, B coordinate represented the color identity of a particular endothelial cell, to measure the clonal relationship among endothelial cells, the distances between each and every endothelial cell were calculated with the formula 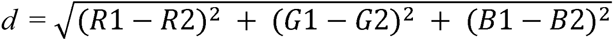. In a matrix, the circle in each cell represented the color distance between two endothelial cells. The distances between clonally related ECs would be small and depicted by small, light blue circles, while the distance between unrelated ECs would be large and indicated by large, dark blue circles. *iii*) 3D RGB intensity scatter plots. RGB values were converted to x, y, z coordinate points on a 3D Cartesian graph. As each individual x, y, z coordinate represented the color identity of a particular endothelial cells, clonally related endothelial cells would be clustered together.

### Tricaine treatment

*Arteriobow;kdrl:mCherry* embryos were treated with 0.028% (2×) tricaine (MS-222, Sigma-Aldrich) from 30 to 54 hpf to slow down heart rate during the secondary sprouting and ISV remodeling. Subsequently, the compound was washed away and embryos re-cultured in E3 embryo medium with 0.003% PTU. At 5 dpf, the identity of arterial or venous ISVs (originally) containing endothelial clones was determined by their connection to, respectively, the DA or the PCV, as well as by direction of blood flow.

### DAPT treatment

*Arteriobow;kdrl:mCherry* embryos were treated with 5 µM DAPT or a similar amount of DMSO, with the latter being used as controls. The treatment window started from 24 hpf until 54 hpf, after which the compound (either DAPT or DMSO) was washed away. The embryos were re-cultured in E3 embryo medium only with 0.003% PTU until 5 dpf to determine the identity of arterial or venous ISVs (originally) containing endothelial clones.

## Acknowledgement

The authors would like to thank Carol Wicking and Holger Gerhardt for helpful discussions, the UQ IMB ACRF imaging facility for technical support.

## Author Contributions

KJ, and MF were involved in the design of the experiments. KJ and CP performed experiments. KJ and ZN were involved in data analyses. MF and KJ wrote the manuscript. KJ and MF were involved in figure construction. All authors commented on the manuscript.

## Data and materials availability

Raw image files are available on Dryad.

---

**Supp. Figure 1 to Figure 1.**
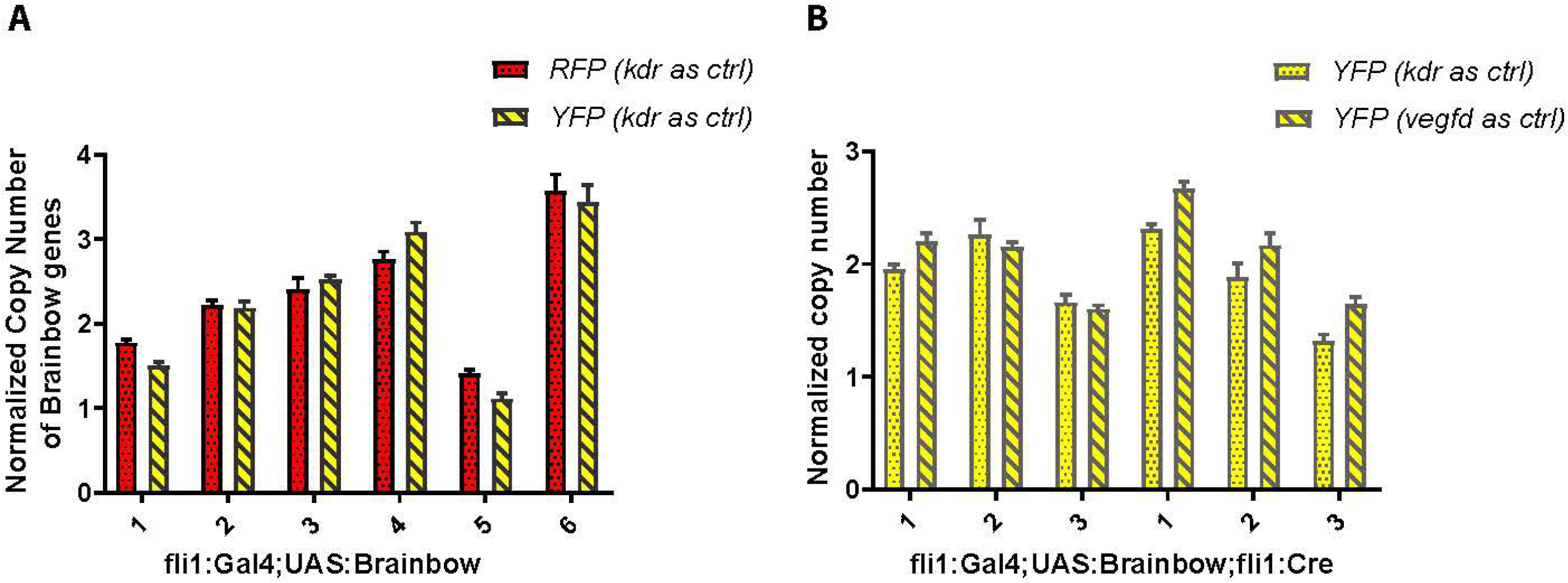
Copy numbers of the Brainbow construct assessed and calculated by qPCR using genomic DNA. (A) qPCR analysis and copy number estimates in genomic DNA from Brainbow transgenic lines without Cre-recombination. kdr (zebrafish gene) and dTomato- and eYFP-coding genes (two Brainbow genes) were used as reference and target genes, respectively, to evaluate the performance of the qPCR method in determining copy numbers of the Brainbow inserts. (B) qPCR analysis of copy numbers in Brainbow transgenic lines with Cre-recombination. dTomato- and mCerulean-coding genes might be excised upon Cre-mediated recombination while eYFP-coding gene always remains in the Brainbow cassette. kdr and vegfd (two zebrafish genes) and eYFP-coding gene were used as reference and target genes, respectively, to evaluate the robustness of the qPCR method in determining copy numbers of the Brainbow inserts. It should be noted that transgenic lines with <= 3 copies of the Brainbow insert(s) were selected as reporters to perform EC cluster analysis in this study.

**Supp. Figure 1 to Figure 2.**
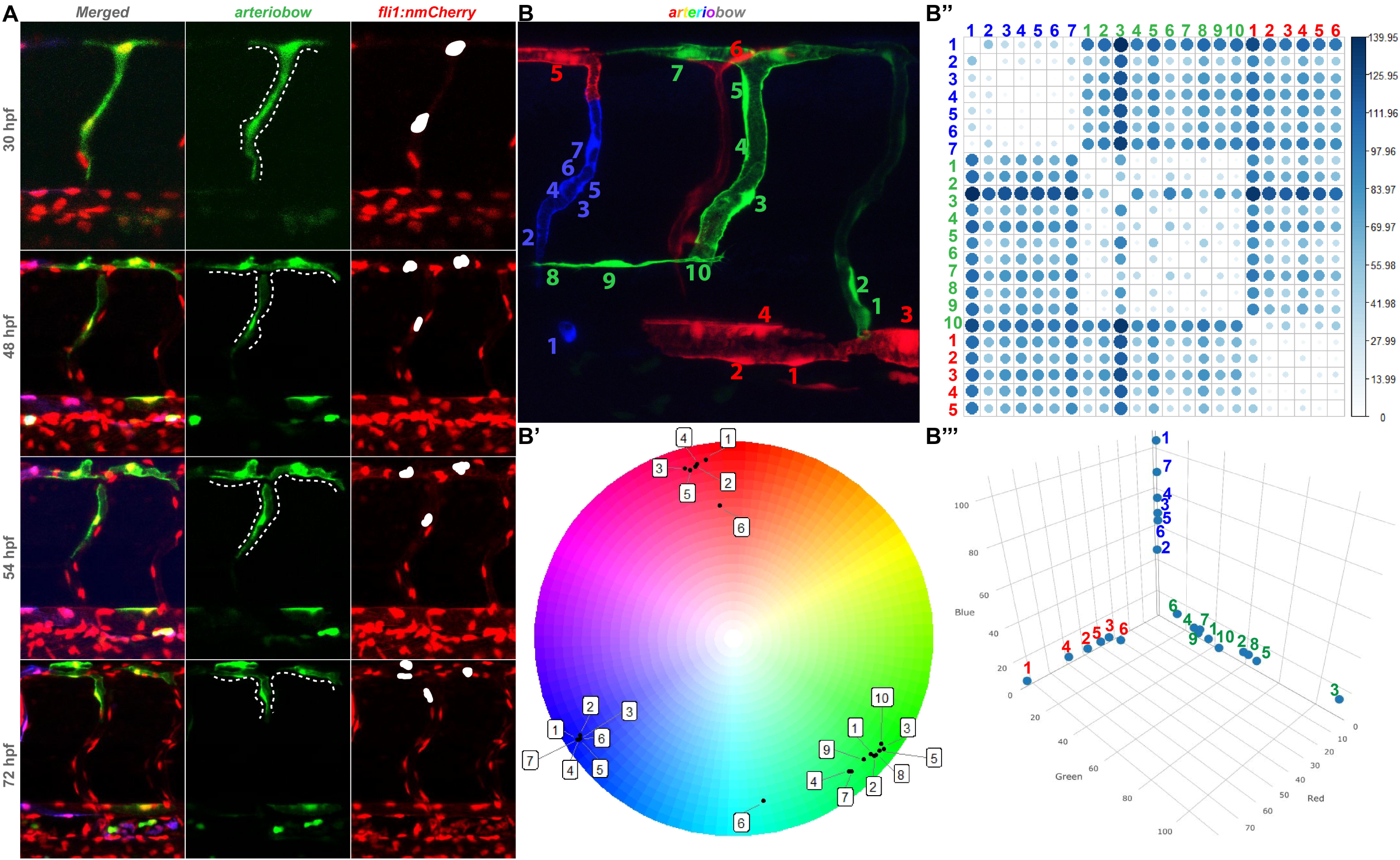
An unbiased approach to measure color distance between cohesive EC clusters in a high throughput manner. **(A) Endothelial cell (EC) nuclei are used as a region of interest to assess fluorescence intensity**. The *arteriobow* line was crossed to the *fli1*:nRFP reporter line where the *fli1* promoter drives expression of nuclear-localized red fluorescence. White staining indicates the nucleus, which is the region of interest within a cell since it is the brightest part of the cell and is subjected to the least amount of fluctuations, in terms of cell shape changes. **(B) Identification of EC nuclei**. The brightest pixels representative of the nuclear area of ECs were automatically selected by the function *find maxima* in Fiji software. The color profiles, RGB intensity values, of the selected ECs were then extracted by running the algorithm “*record maxima colors*” (each number indicates a nucleus). **(B’) Positioning colored EC clusters on a color wheel using hue and saturation information**. RGB intensity values of the selected EC nuclei are converted into hue and saturation values. EC clusters were plotted onto a color wheel where the hue value represents the color information of the cluster, while saturation represents the intensity of the color. The relative position of the cluster from the center of the wheel is proportional to color intensity, with low intensity close to center and high intensity at the periphery. **(B’’) Visualizing color distances with a matrix to define relationship**. The color distance is calculated using 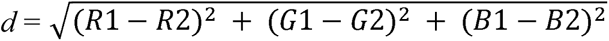. The distance between closely related color is depicted by small light blue circles, while the distance between unrelated color is shown by a large dark blue circle. **(B’’’) Defining color distance on a 3D RGB intensity scatter plot**. RGB intensity values were converted to x, y, z co-ordinate points on a 3D Cartesian graph. Given that each individual co-ordinate represents the color identity of a specific EC, ECs from the same cohort with closely related colors assemble in clusters. Relationship between ECs within the same cluster and among different clusters is resolved by combining the three calculations shown in B’, B” and B’’’. hpf, hours post fertilization.

**Supp. Figure 2 to Figure 2.**
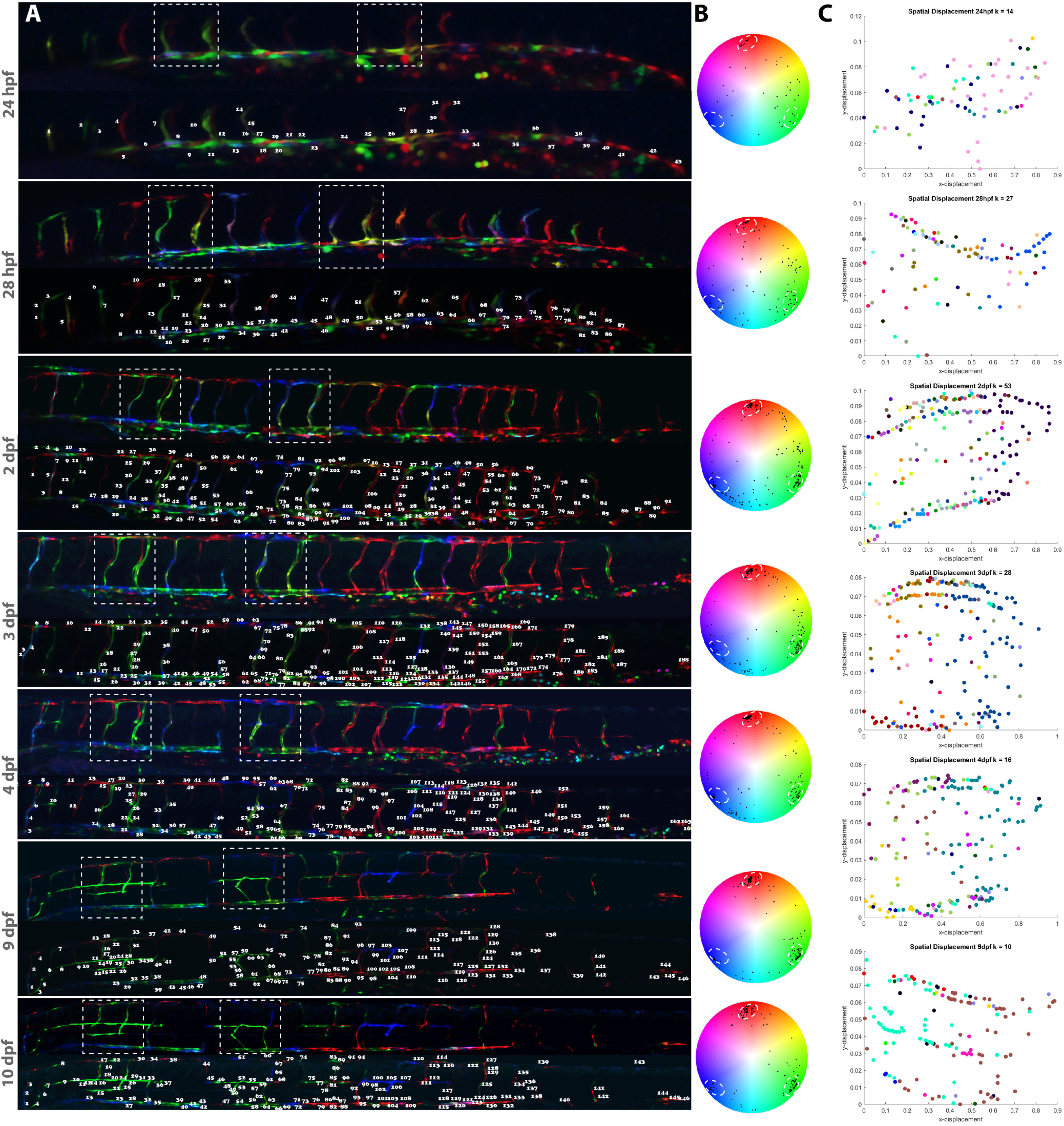
Heterogeneity in collective EC dynamics is conserved over time in the developing blood vasculature. (**A**) Whole trunk vasculature imaged over time from 24 hpf until 10 dpf using confocal microscopy. Unbiased selection of nuclei from all ECs labelled by the Brainbow cassette and annotated individually with a specific number. (**B**) Clonal clustering analysis based on color distance. (**C**) Clustering analysis performed over time using K-means methods. It should be noted that the color coding at each developmental stage is unique to itself, whereby each color represents an individual clone at that developmental stage only. hpf, hours post fertilization; dpf, days post fertilization.

**Supp. Figure 1 to Figure 3.**
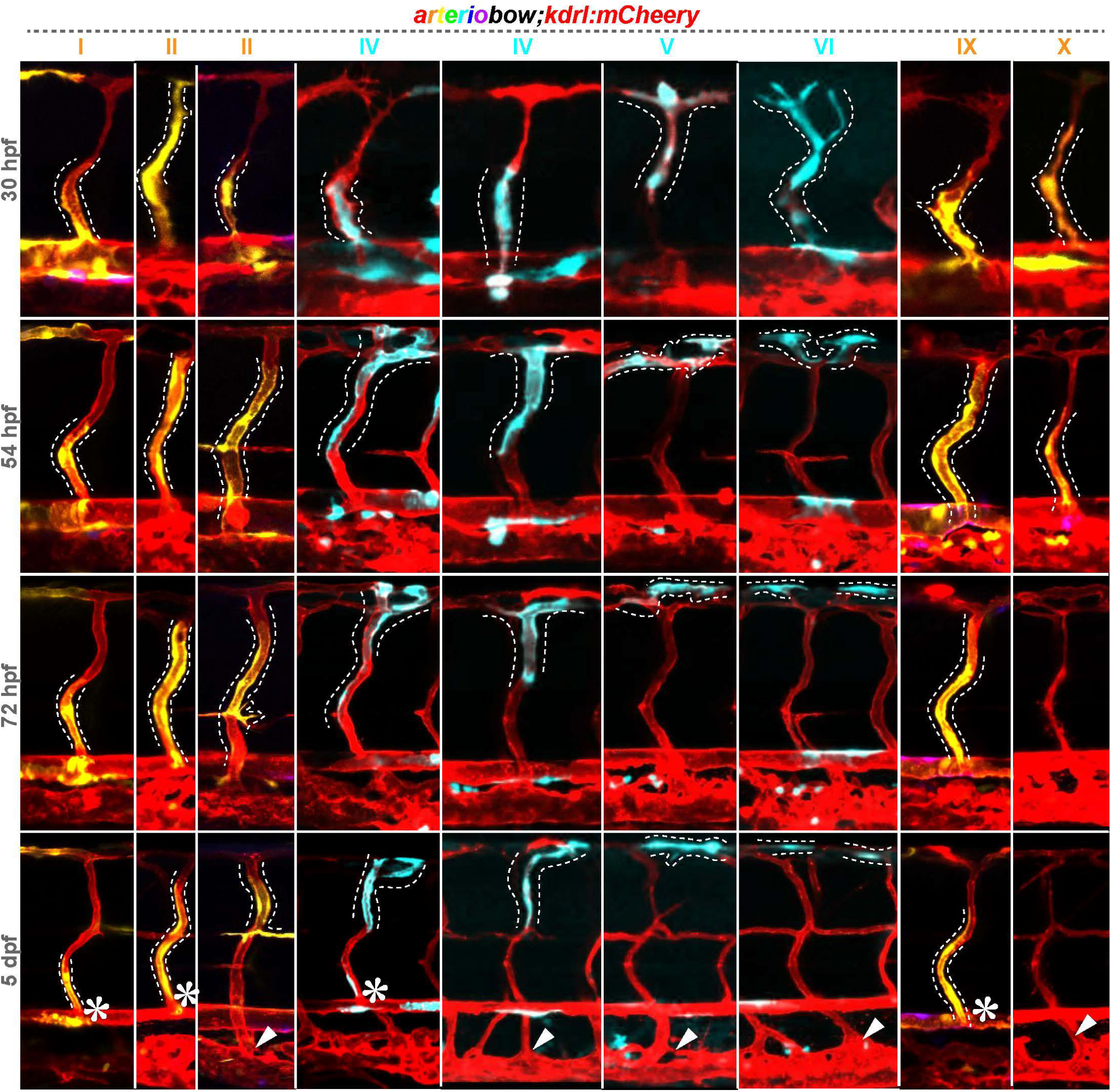
Map of collective behaviors of arterial ECs in relation to arterio-venous remodeling. The *arteriobow* line was crossed to the *kdrl:mCherry* line and behavioral types of EC cohorts were analyzed and mapped in relation to arterio-venous remodeling. Representative collective EC behaviors (I, II, IV, V, VI, IX and X) are shown by a white dashed line. Asterisks denote the connection between aISVs and the DA, while arrowheads show the connection between vISVs and the PCV. hpf, hours post fertilization; dpf, days post fertilization; aISV, arterial intersegmental vessel; DA, dorsal aorta; DLAV, dorsal longitudinal anastomosing vessel.

**Supp. Figure 2 to Figure 3.**
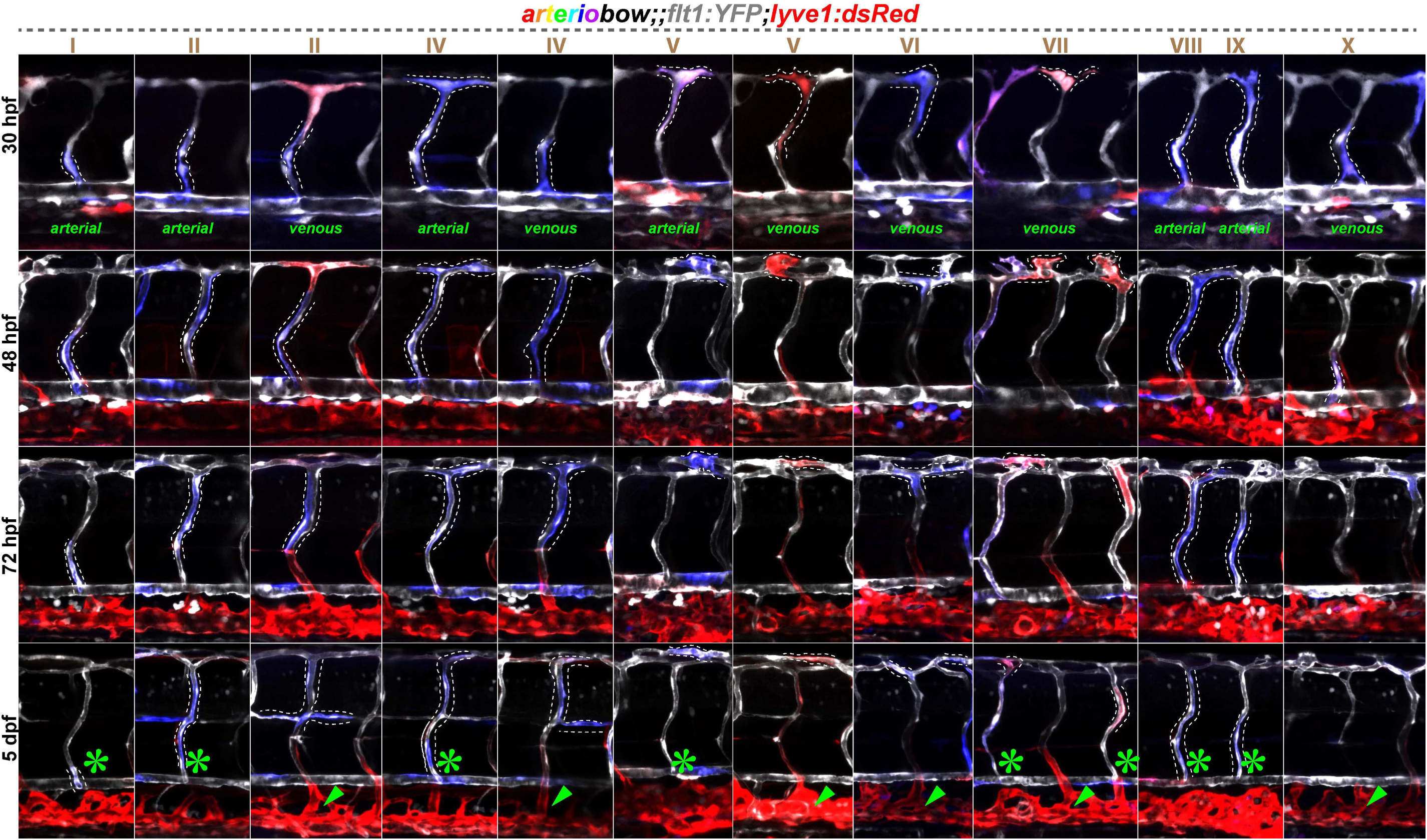
Verification of differential behavior of cohesive EC clusters as a driver in arterio-venous remodeling. The *arteriobow* line was crossed to the *flt1:YFP;lyve1:dsRed* line and collective EC behaviors were analyzed and mapped in relation to arterio-venous remodeling. Behavioral types are outlined by white dashed lines. Asterisks denoted the connection between aISVs and the DA, while arrowheads showed the connection between vISVs and the PCV. hpf, hours post fertilization; dpf, days post fertilization; aISV, arterial intersegmental vessel; DA, dorsal aorta; DLAV, dorsal longitudinal anastomosing vessel.

**Supp. Figure 3 to Figure 3.**
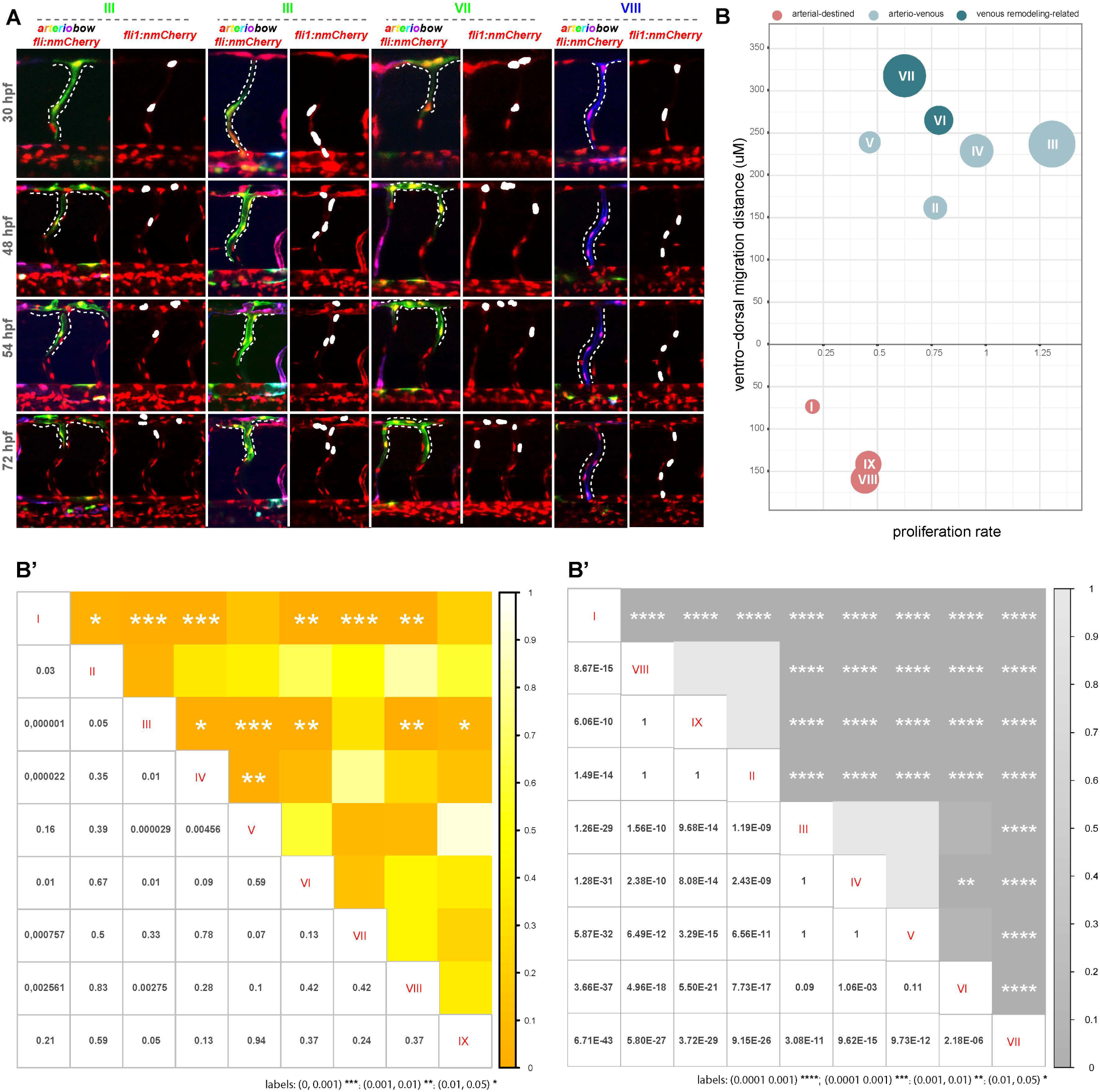
Quantification of cluster size and proliferation rate. (**A**) The *arteriobow* line was crossed to the *fli1:nmCherry* line to analyze the proliferative behavior of cohesive EC cohorts over a 42 h period (30hpf to 72hpf). Representative clonal types at four different time points during vascular remodeling are shown. The type III clones are representative of endothelial clones capable of dorsal migration; the type VII rearranges during vascular remodeling; the type VIII represent actively proliferating endothelial clones with a restricted remodeling specific to aISV. Clones of interest are outlined by white dashed lines and nuclei within each clone are highlighted in white. (**B**-**B**’’) Heterogeneity in the migratory and proliferative behaviors of cohesive EC clusters (n = 6 experiments, 33 embryos, 235 clones). (**B**) Proliferation rate, migration distance, and cluster size of different types of collective EC cohorts. The x-axis represents the mean proliferation rate, which is determined by using the formula s = (N_2_ – N_1_)/42, where N_1_ and N_2_ represent the number of endothelial nuclei within a cluster at 30 hpf and 72 hpf, respectively. The average migration distance of cohesive EC clusters is shown on the y-axis. It is quantified at 72 hpf by measuring the distance between the front end of EC cohorts and the DA. The radius of the bubble is proportional to the cluster size, which is acquired by counting the average number of nuclei within each behavioral type at 72 hpf. (**B’-B’’**) Matrix heatmaps of p-values obtained from performing pairwise t-tests for proliferation rate (B’) and migration distance (B’’) of multiple types of collective EC behaviors. The lower triangular matrix shows the p-values obtained from t-test, and the upper triangular matrix shows significance levels. The t-test as implemented in R was used for statistical analysis; significance (*P < 0.05, **P < 0.01, ***P < 0.001, ****P < 0.0001) in unpaired t-test; not significant if not marked by a significance label). hpf, hours post fertilization.

**Supp. Figure 1 to Figure 4.**
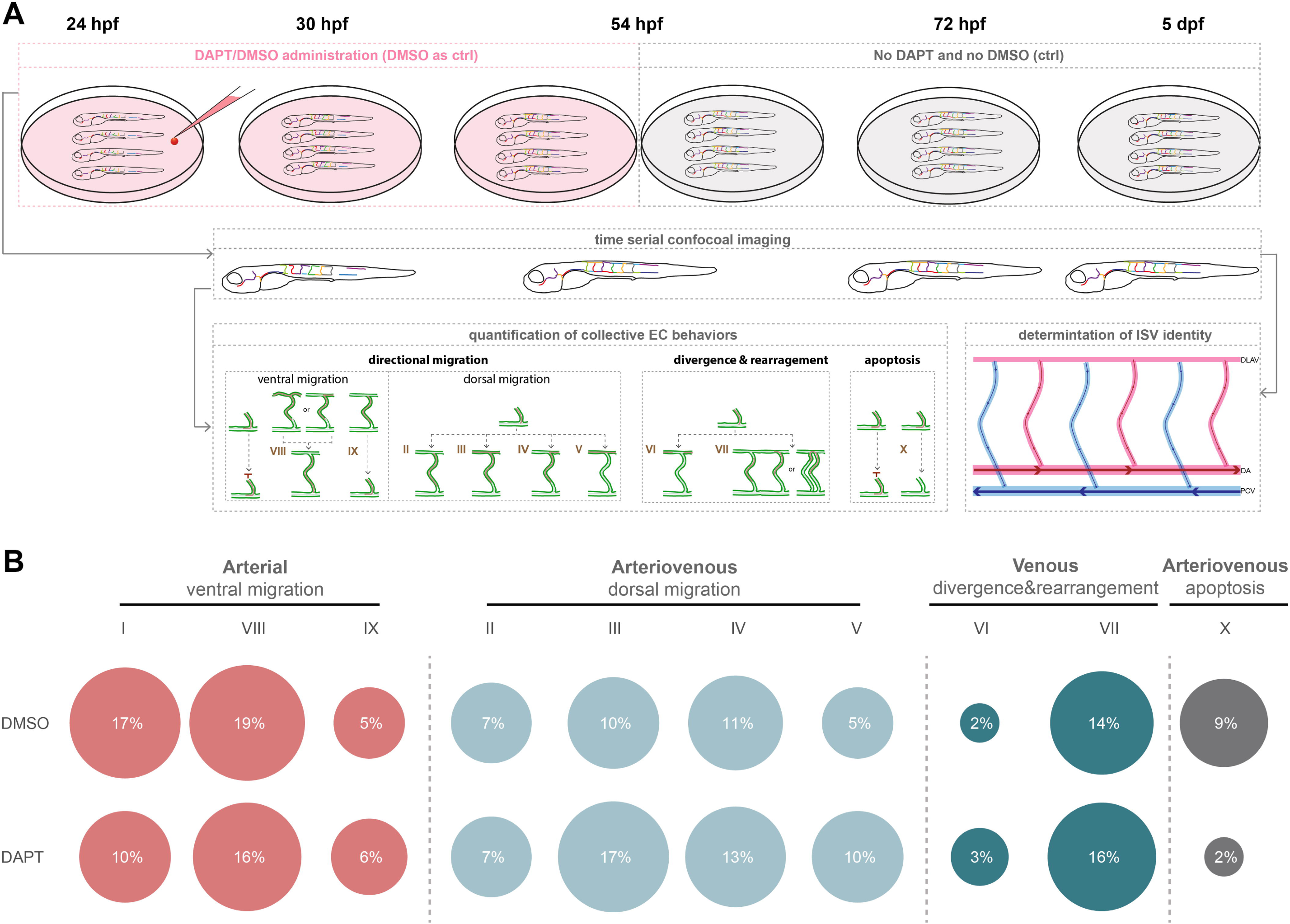
**(A)** Workflow for manipulating Notch signaling to study its effects on collective EC behaviors. *arteriobow;kdrl:mCherry* embryos were treated with DAPT at 24 hpf. Time-course confocal imaging was performed across different developmental stages, 30 hpf, 54 hpf, 72 hpf and 5 dpf, respectively. Ten types of collective EC behaviors were characterized and quantified. The identity of ISVs was determined at 5 dpf by studying their connection with either the DA or PCV, as well as the direction of blood flow. **(B)** Quantification of the proportion of cohesive EC clusters in each of the 10 behavior classes in the *arteriobow;kdrl:mCherry* embryos either untreated or treated with DAPT. The top panel shows the proportion of the ten behavioral types in basal conditions, the bottom shows the behavioral distribution in the presence of DAPT. hpf, hours post fertilization; dpf, days post fertilization; aISV, arterial intersegmental vessel; DA, dorsal aorta; DLAV, dorsal longitudinal anastomosing vessel.

**Supp. Figure 1 to Figure 5.**
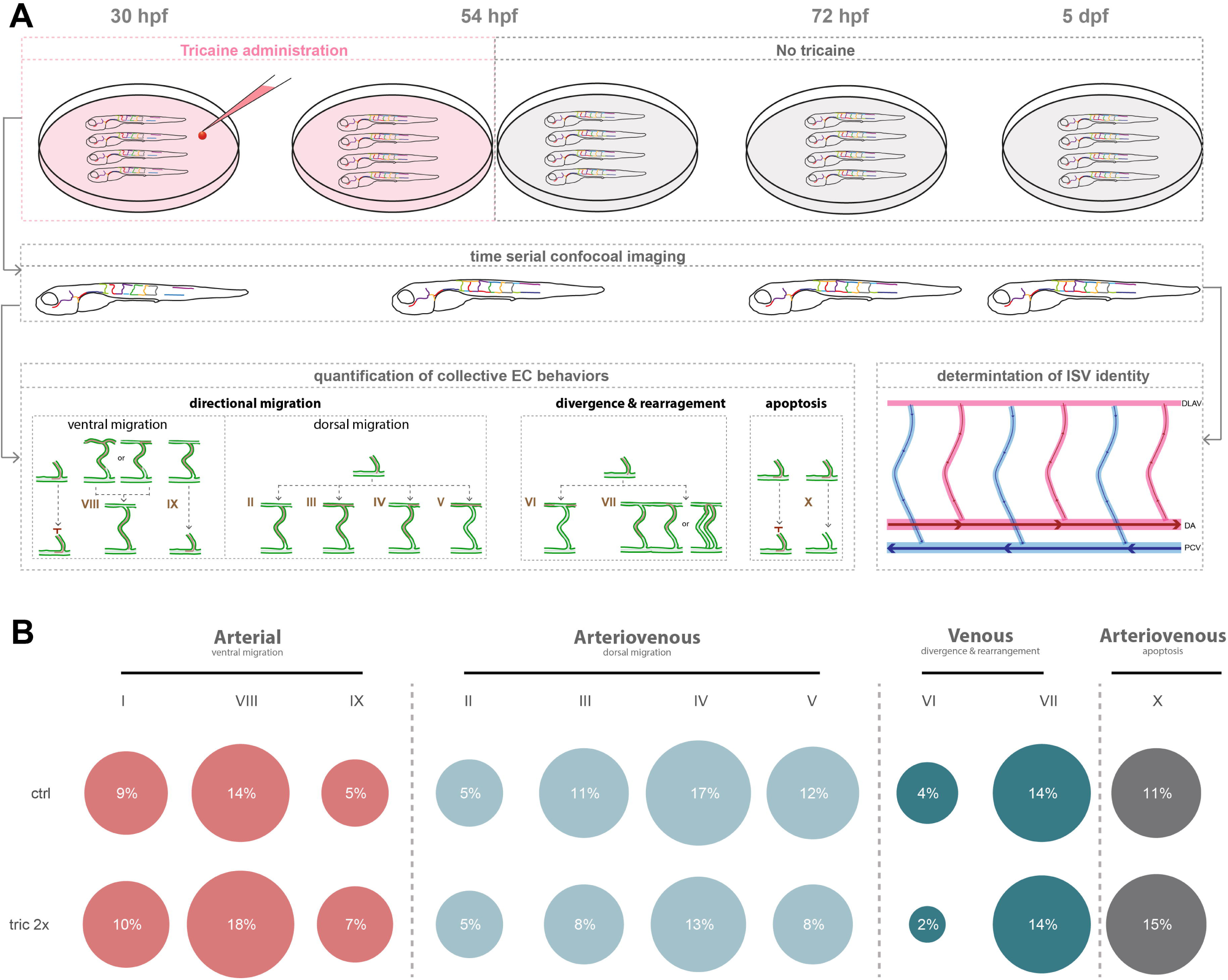
**(A)** Workflow for manipulating blood flow to study its effects on collective EC behaviors. *arteriobow;kdrl:mCherry* embryos were treated with 2× tricaine to slow down the heart rate and blood flow from 30 hpf to 54 hpf. Time-course confocal imaging was performed across different developmental stages, 30 hpf, 54 hpf, 72 hpf and 5 dpf, respectively. 10 types of collective EC behaviors were characterized and quantified. The identity of ISVs was determined at 5 dpf by studying their connection with either the DA or PCV, as well as the direction of blood flow. **(B)** Quantification of the proportion of cohesive EC clusters in each of the 10 behavioral types was performed in the trunk of 5 dpf *arteriobow;kdrl:mCherry* embryos either untreated (control; n=6 experiments, 42 embryos, 333 clusters) or treated with tricaine (n=6 experiments, 42 embryos, 471 clusters) to slow down blood flow. The top panel represents the percentage of EC clusters of each behavioural type in basal conditions, the bottom panel shows the distribution of collective EC behaviors in the presence of tricaine. hpf, hours post fertilization; dpf, days post fertilization; aISV, arterial intersegmental vessel; DA, dorsal aorta; DLAV, dorsal longitudinal anastomosing vessel.

## References

Aird, W. C. (2012). “Endothelial cell heterogeneity.” Cold Spring Harbor perspectives in medicine 2(1): a006429.

Asakawa, K. and K. Kawakami (2008). “Targeted gene expression by the Gal4-UAS system in zebrafish.” Dev Growth Differ 50(6): 391–399.

Bussmann, J., et al. (2010). “Arteries provide essential guidance cues for lymphatic endothelial cells in the zebrafish trunk.” Development 137(16): 2653–2657.

Chen, Q., et al. (2012). “Haemodynamics-driven developmental pruning of brain vasculature in zebrafish.” PLoS Biol 10(8): e1001374.

Chen, Q., et al. (2012). “Haemodynamics-driven developmental pruning of brain vasculature in zebrafish.” PLoS biology 10(8).

Chiang, I. K.-N., et al. (2017). “SoxF factors induce Notch1 expression via direct transcriptional regulation during early arterial development.” Development 144(14): 2629–2639.

Dahlem, T. J., et al. (2012). “Simple Methods for Generating and Detecting Locus-Specific Mutations Induced with TALENs in the Zebrafish Genome.” PLOS Genetics 8(8): e1002861.

Fang, J. S., et al. (2017). “Shear-induced Notch-Cx37-p27 axis arrests endothelial cell cycle to enable arterial specification.” Nature Communications 8(1): 1–14.

Franco, C. A., et al. (2015). “Dynamic endothelial cell rearrangements drive developmental vessel regression.” PLoS biology 13(4).

Geudens, I., et al. (2019). “Artery-vein specification in the zebrafish trunk is pre-patterned by heterogeneous Notch activity and balanced by flow-mediated fine-tuning.” Development 146(16): dev181024.

Geudens, I. and H. Gerhardt (2011). “Coordinating cell behaviour during blood vessel formation.” Development 138(21): 4569–4583.

Geudens, I., et al. (2010). “Role of delta-like-4/Notch in the formation and wiring of the lymphatic network in zebrafish.” Arteriosclerosis, thrombosis, and vascular biology 30(9): 1695–1702.

Herbert, S. P. and D. Y. Stainier (2011). “Molecular control of endothelial cell behaviour during blood vessel morphogenesis.” Nature Reviews Molecular Cell Biology 12(9): 551–564.

Hogan, B. M., et al. (2009). “Vegfc/Flt4 signalling is suppressed by Dll4 in developing zebrafish intersegmental arteries.” Development 136(23): 4001–4009.

Hogan, B. M. and S. Schulte-Merker (2017). “How to plumb a pisces: understanding vascular development and disease using zebrafish embryos.” Developmental Cell 42(6): 567–583.

Hughes, S. and T. Chan-Ling (2000). “Roles of endothelial cell migration and apoptosis in vascular remodeling during development of the central nervous system.” Microcirculation 7(5): 317–333.

Isogai, S., et al. (2003). “Angiogenic network formation in the developing vertebrate trunk.” Development 130(21): 5281–5290.

Jakobsson, L., et al. (2010). “Endothelial cells dynamically compete for the tip cell position during angiogenic sprouting.” Nature cell biology 12(10): 943–953.

Jianxin, A. Y., et al. (2015). “Single-cell analysis of endothelial morphogenesis in vivo.” Development 142(17): 2951–2961.

Kimmel, C. B., et al. (1995). “Stages of embryonic development of the zebrafish.” Developmental dynamics 203(3): 253–310.

Kohli, V., et al. (2013). “Arterial and venous progenitors of the major axial vessels originate at distinct locations.” Developmental Cell 25(2): 196–206.

Küchler, A. M., et al. (2006). “Development of the zebrafish lymphatic system requires VEGFC signaling.” Current Biology 16(12): 1244–1248.

Ladoux, B. and R.-M. Mège (2017). “Mechanobiology of collective cell behaviours.” Nature Reviews Molecular Cell Biology 18(12): 743–757.

Lawson, N. D., et al. (2001). “Notch signaling is required for arterial-venous differentiation during embryonic vascular development.” Development 128(19): 3675–3683.

Le Noble, F., et al. (2004). “Flow regulates arterial-venous differentiation in the chick embryo yolk sac.” Development 131(2): 361–375.

Liu, F., et al. (2008). “Fli1 acts at the top of the transcriptional network driving blood and endothelial development.” Curr Biol 18(16): 1234–1240.

Livet, J., et al. (2007). “Transgenic strategies for combinatorial expression of fluorescent proteins in the nervous system.” Nature 450: 56.

Lobov, I. B., et al. (2005). “WNT7b mediates macrophage-induced programmed cell death in patterning of the vasculature.” Nature 437(7057): 417–421.

Lucitti, J. L., et al. (2007). “Vascular remodeling of the mouse yolk sac requires hemodynamic force.” Development 134(18): 3317–3326.

Mishra, A. K., et al. (2019). “Cell interactions in collective cell migration.” Development 146(23).

Nguyen, P. D., et al. (2017). “Muscle stem cells undergo extensive clonal drift during tissue growth via Meox1-mediated induction of G2 cell-cycle arrest.” Cell Stem Cell 21(1): 107-119. e106.

Pan, Y. A., et al. (2011). “Multicolor Brainbow imaging in zebrafish.” Cold Spring Harb Protoc 2011(1): pdb.prot5546.

Park, C., et al. (2013). “Transcriptional regulation of endothelial cell and vascular development.” Circ Res 112(10): 1380–1400.

Pichol-Thievend, C., et al. (2018). “A blood capillary plexus-derived population of progenitor cells contributes to genesis of the dermal lymphatic vasculature during embryonic development.” Development 145(10).

Potente, M. and T. Mäkinen (2017). “Vascular heterogeneity and specialization in development and disease.” Nature Reviews Molecular Cell Biology 18(8): 477.

Rochon, E. R., et al. (2016). “Alk1 controls arterial endothelial cell migration in lumenized vessels.” Development 143(14): 2593–2602.

Scarpa, E. and R. Mayor (2016). “Collective cell migration in development.” Journal of Cell Biology 212(2): 143–155.

Schindelin, J., et al. (2012). “Fiji: an open-source platform for biological-image analysis.” Nature methods 9(7): 676–682.

Scott, E. K., et al. (2007). “Targeting neural circuitry in zebrafish using GAL4 enhancer trapping.” Nature Methods 4: 323.

Su, T., et al. (2018). “Single-cell analysis of early progenitor cells that build coronary arteries.” Nature 559(7714): 356–362.

Sumanas, S. and S. Lin (2005). “Ets1-related protein is a key regulator of vasculogenesis in zebrafish.” PLoS Biol 4(1): e10.

Tatapudy, S., et al. (2017). “Cell fate decisions: emerging roles for metabolic signals and cell morphology.” EMBO reports 18(12): 2105–2118.

Thüroff, F., et al. (2019). “Bridging the gap between single-cell migration and collective dynamics.” Elife 8: e46842.

Torres-Vázquez, J., et al. (2003). “Molecular distinction between arteries and veins.” Cell and tissue research 314(1): 43–59.

Udan, R. S., et al. (2013). “Dynamic responses of endothelial cells to changes in blood flow during vascular remodeling of the mouse yolk sac.” Development 140(19): 4041–4050.

Wacker, A. and H. Gerhardt (2011). “Endothelial development taking shape.” Current opinion in cell biology 23(6): 676–685.

Weijts, B., et al. (2018). “Blood flow-induced Notch activation and endothelial migration enable vascular remodeling in zebrafish embryos.” Nature Communications 9(1): 1–11.

Xu, C., et al. (2014). “Arteries are formed by vein-derived endothelial tip cells.” Nature Communications 5(1): 1–11.

Yaniv, K., et al. (2006). “Live imaging of lymphatic development in the zebrafish.” Nature medicine 12(6): 711–716.

Zhong, T. P., et al. (2001). “Gridlock signalling pathway fashions the first embryonic artery.” Nature 414(6860): 216–220.

Zhong, T. P., et al. (2000). “gridlock, an HLH gene required for assembly of the aorta in zebrafish.” Science 287(5459): 1820–1824.

